# Clinical and Molecular Characterization of Glioma Patients: miR-21 Expression as a Prognostic Biomarker in Tissue and Serum

**DOI:** 10.1101/2024.11.20.624501

**Authors:** Altaf Ali Laghari, Sufiyan Sufiyan, Wajiha Amin, Umer Adnan, Sana Naeem, Syed Hani Abidi, Sahar Ilyas, Siraj Uddin, Mohammad Hamza Bajwa, Syed Ather Enam, Nouman Mughal

**Affiliations:** Department of Surgery, Aga Khan University Hospital, Karachi, Pakistan; Department of Biological & Biomedical Science, Aga Khan University Hospital, Karachi; Department of Biomedical Sciences, Nazarbayev School of Medicine, Nazarbayev University, Nur-Sultan, Kazakhstan; Center of Oncological Research in Surgery, Aga Khan University; Medical College Aga Khan University, Karachi, Pakistan

**Keywords:** Glioma, Biomarker, Molecular markers, miRs, Liquid biopsy

## Abstract

**Background:** Glioma remains challenging due to high recurrence rates and resistance to treatment. Diagnosis and follow-up in resource-constrained regions often leads to significant patient attrition. Serum microRNA (miRNA) expression profiles, which have been shown to correlate with tissue expression profiles, are detectable in peripheral blood samples, providing a promising avenue for non-invasive and repeatable liquid biopsies. miR-21 shows promise in many populations; however, there is a dearth of data from our region.

**Methodology:** We collected 90 tumor tissues, 42 pre- and post-operative serum samples from glioma patients, and included 10 normal tissue adjacent to the tumor (NATs) along with serum samples from 8 healthy individuals and analyzed for miR-21 expression through RT-qPCR. Shapiro Wilk test was applied to calculate data distribution, ANOVA, Fisher’s exact, and Wilcox test, along with pairwise Student’s t-test, were applied to determine the differences in gene expression. The expression level of miR-21 was assessed for correlation with Kaplan-Meier survival curves and different molecular markers (IDH, Ki-67, ATRX and p53). The quantitative hazard ratio was determined using Cox regression analysis.

**Results:** miR-21 expression in tissue increased with the mean log fold expression 0.101 (median fold = 2.35) in grade 2, 1.00 (median fold = 7.49) in grade 3 and 1.53 (median fold = 26.0) in grade 4 for glioma patients. The expression level showed significant difference between control tissue and grade 4 patients along with significant inter-comparison between grade 1 and grade 4, as well as grade 2 and grade 4.A significant elevated expression of miR-21 has been noticed in patients above 50 years of age. Similarly, in serum samples a significant decline in miR-21 expression was observed in post-operative samples as compared to pre-operative samples mean log fold in grade 2 is 1.30 (11.6-fold), grade 3 is 1.08 (15.3-fold) and grade 4 is 0.749 (13.2-fold). Furthermore, there was positive correlation of miR-21 expression with tumor volume. IDH-wildtype and high Ki-67 expression in gliomas showed significant upregulation of miR-21 compared to IDH-mutant and low Ki-67 respectively. Patients with low miR-21 expression had significantly longer overall survival (OS) than patients with high miR-21 expression. Quantitative hazard analysis indicates that patients in the high expression group have a 3.4 times higher risk of mortality (95% CI: 1.6-7.1), in comparison to patients in the low expression group with AUC of 0.742 (all p <0.05).

**Conclusion:** This study demonstrates the potential of microRNA 21 as a serum biomarker for early, cost-effective diagnosis of glioma. Furthermore, it may inform the development of targeted treatment strategies for various glioma grades, particularly in our population.

## Introduction

Gliomas represent the most prevalent primary central nervous system (CNS) malignancies, accounting for approximately 30% of all CNS tumors and 80% of malignant brain tumors [1]. According to the 2021 World Health Organization (WHO) classification, gliomas are categorized into four grades (grades 1-4) based on histological characteristics, microscopic features, and molecular profiles [2]. Despite significant advancements in surgical techniques and adjuvant therapies, the median overall survival for glioblastoma (GBM), the most aggressive form of glioma (grade 4), remains poor. In developed countries, the median survival for GBM patients ranges from 12 to 15 months, primarily due to access to state-of-the-art medical facilities, advanced therapeutic interventions, and comprehensive post-treatment care [3]. In contrast, low-and-middle-income countries (LMICs) report significantly lower median survival rates, ranging from 6 to 9 months. This disparity primarily stems from limited infrastructure and access to advanced diagnostic tools as well as higher rates of treatment discontinuation due to financial and logistical constraints. The prognosis is further worsened by the inherent complexity of treating malignant brain tumors, which necessitates expensive imaging scans, prolonged treatment regimens, and intensive post-surgical follow-ups [4]. These challenges often lead to significant patient dropout rates, underscoring the urgent need for more accessible and cost-effective diagnostic and prognostic tools.

Given these constraints, non-invasive serum-based liquid biopsies are emerging as practical alternatives for glioma diagnosis and monitoring across all grades [5]. Among various candidates, microRNAs (miRs), which are small non-coding RNAs approximately 18-22 nucleotides in length, have shown considerable promise. MiRs regulate gene expression at the post-transcriptional level and are involved in essential biological processes, including cell proliferation, differentiation, apoptosis, and angiogenesis [6, 7]. Extracellular miRs are notably stable and detectable in various body fluids such as serum, plasma, urine, and cerebrospinal fluid (CSF), making them ideal for use as biomarkers [8].

One such miRNA, miR-21, is a well-established oncogenic miRNA implicated in various malignancies, including both solid tumors (e.g., liver, lung, esophageal, and head and neck cancers) and non-solid tumors (e.g., chronic lymphocytic leukemia and B cell lymphoma) [9, 10]. The overexpression of miR-21 has been consistently linked to the promotion of tumor growth, invasion, and metastasis, making it a critical player in cancer biology. In the context of gliomas, miR-21 is markedly upregulated and exerts its oncogenic effects by downregulating several key tumor suppressor genes, such as PTEN (Phosphatase and Tensin Homolog) and PDCD4 (Programmed Cell Death Protein 4) [11]. This miRNA modulates essential oncogenic pathways, including the Wnt/β-catenin signaling pathway, which is crucial for cell proliferation and differentiation, and the STAT3 (Signal Transducer and Activator of Transcription 3) pathway, which plays a significant role in tumorigenesis and immune evasion [12]. Additionally, miR-21 impacts the PI3K/AKT/mTOR pathway, a central regulator of cell survival, growth, and metabolism, further contributing to the aggressive nature of gliomas [13]. Studies have demonstrated that the overexpression of miR-21 correlates with higher tumor grades, enhanced proliferative capacity, and reduced survival rates in glioma patients, indicating its potential as a prognostic marker. A comprehensive understanding of miR-21’s involvement in these processes highlights its value as a biomarker for both the diagnosis and prognosis of gliomas.

Despite the established role of miR-21 in glioma biology, there is a significant gap in data regarding its relevance and clinical utility in specific populations, particularly in LMICs. Addressing this gap is crucial, as population-specific genetic and environmental factors may influence miR-21 expression, leading to variations in the diagnostic performance and prognostic utility as well as therapeutic response [14]. Additionally, the data collected in this study is essential for guiding future research aimed at describing the role of miR-21 in pathogenesis of glioma.

To address these gaps, we designed a study to evaluate the potential of circulating miR-21 as a non-invasive biomarker in both tissue and serum samples from glioma patients treated at a tertiary-care hospital. Our objectives include assessing miR-21 expression levels and their correlations with clinical parameters, tumor grade (1-4), the association of miR-21 levels in tissue and serum with patient overall survival (OS) to determine its prognostic value. Furthermore, we will evaluate the specificity of miR-21 as a biomarker by analyzing its performance in distinguishing between glioma patients and controls, using the Area Under the Curve (AUC) of Receiver Operating Characteristic (ROC) curves. Furthermore, enrichment analysis was performed using the miR database (MiRDB) to identify miR-21 target genes and associated pathways, aiming to explore some of the molecular mechanisms underlying miR-21’s role in glioma genesis.

## Methodology

### Study Design

This study was approved by the Institutional Ethical Review Committee (AKU-ERC # 2019-1945-5110, AKU-ERC # 2021-1945-17282; AKU-ERC # 2021-6265-18766) at Aga Khan University Hospital, Karachi, Pakistan. Written informed consent was obtained from all participants prior to surgery. Patients diagnosed with glioma based on final histopathology reports were recruited. Patient data and medical histories were collected from the hospital electronic medical records. Patients were followed up to extract survival data, with the follow-up period defined from the date of surgery until the patient’s death or the last 5 years of follow-up. A written informed consent was also taken from healthy individuals whose serum samples were used as controls.

### Sample Collection

A total of 90 glioma tumor tissues, along with 10 adjacent non-tumor tissues, were collected and stored at − 80 °C at the time of surgery. Serum aliquots were prepared from blood samples obtained from 42 patients before and 24 hours after surgery and stored under the same conditions. Additionally, serum samples from 8 healthy individuals were collected for control analysis. The discrepancy in sample numbers between tissue and serum samples arose due to various challenges. For some patients, blood samples were compromised before serum separation, and replacements were impractical post-surgery. In other cases, serum samples had compromised RNA quality and quantity due to mishandling during storage or inadequate conditions. Some samples yielded insufficient RNA, resulting in elevated cycle threshold (CT) values for housekeeping genes, indicating minimal gene expression.

### Immunohistochemistry of Molecular Markers

Histological samples of glioma formalin-fixed paraffin-embedded (FFPE) tissues were examined and graded by a histopathologist at Aga Khan University Hospital. Molecular marker analysis, including IDH (wildtype/mutant), ATRX (loss/retained), Ki67 (low/high), and p53 (wildtype/mutant), was conducted using immunohistochemical (IHC) staining. Patient data was retrieved from hospital medical records for comprehensive analysis.

### RNA Extraction

Total RNA was isolated from tumor tissue samples using the Trizol chloroform method. RNA extraction from 200 μl of serum was performed with the miRNeasy Serum/Plasma Kit (Qiagen). The quantity of isolated RNA was assessed using NanoDrop™ 2000/2000c Spectrophotometers (Thermo Scientific). To remove DNA contamination, RNA samples underwent DNase I treatment using the DNA-free DNase Treatment Kit (Thermo Fisher) following the manufacturer’s protocol.

### cDNA Synthesis

To assess miR-21 expression levels in tissue and serum samples, 1 μg of RNA was reverse transcribed using the RevertAid First Strand cDNA Synthesis Kit (Thermo Fisher). A reaction mixture of 5X reaction buffer, 1μl Ribolock inhibitor (RI), 1 μl RevertAid reverse transcriptase enzyme (RT), 2 μl dNTPs (10 μM), 1ug treated RNA and nuclease free water to make up the volume up to 20μl. The reaction was subjected to temperature cycles of 42 °C for 60 minutes and 70°C for 5 minutes. The prepared cDNA product was stored at −20 °C until further use. During cDNA synthesis for miR-21, a stem-loop (SLRT) primer specific to miR-21 sequences was employed to enhance specificity. U6 expression levels were established using a specific SLRT primer during reverse transcription. U6 expression served as housekeeping genes to normalize miR-21 expression in both tissue and serum samples. For both miR-21 and U6, qPCR was performed with specific forward and reverse primers. The sequences of the SLRT primer and the specific primers for miR-21 and U6 are mentioned in Table 1.

**Table 1:**
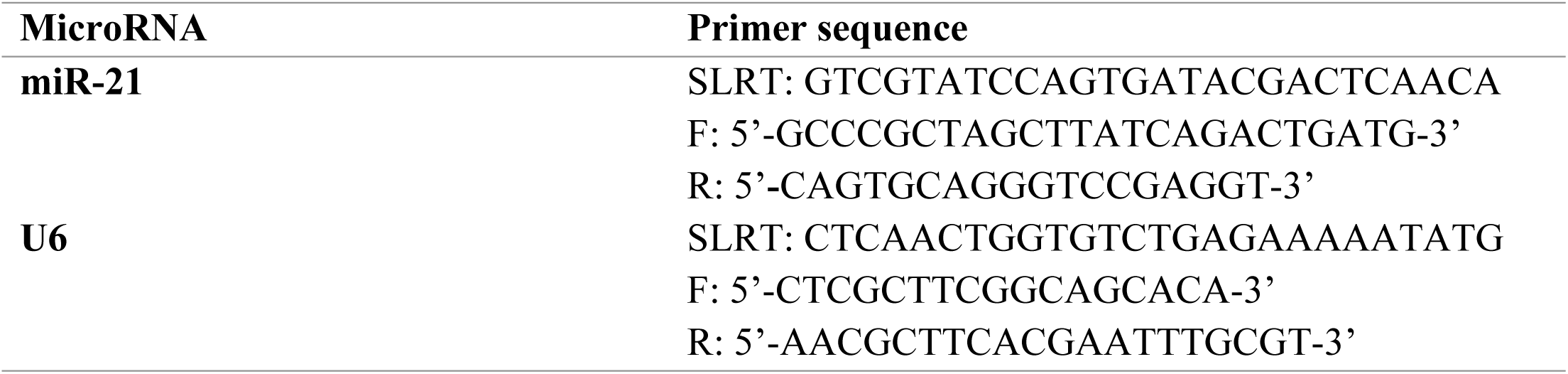
The list of primer sequences used in this study.

### miR-21 Quantitative Analysis by q-PCR

For the assessment of miR-21 expression, qPCR was performed using a reaction mix of 10 µl, which comprised 2 μl of cDNA template, 5 μl of PowerUp™ SYBR™ Green Master Mix (Thermo Fisher Scientific), 1 μl each of forward and reverse primers (10 μM) (Eurofins, USA), and 2 μl of nuclease-free water. The thermal cycling conditions included an initial hold for 2 minutes at 50°C, followed by another hold for 2 minutes at 95°C, and then 40 cycles of denaturation at 95°C for 15 seconds and annealing/extension at 60°C for 60 seconds. Each reaction was conducted in duplicate, with non-template control (NTC) and non-primer control (NPC) included as experimental controls. The U6 gene, a small nuclear RNA gene, was utilized as an endogenous control. The relative fold change in miR-21 expression in each sample was calculated using the Livak 2^−ΔΔCt^ method [15].

### Statistical Analysis

The hypothesis test for data distribution was calculated using the Shapiro-Wilk test. ANOVA and Fisher’s exact test, along with pairwise Student’s t-test, were applied to determine the differences in gene expression between the grades. Pearson correlation was used to determine the association pattern of tumor volume with miR-21 expression. The expression difference of miR-21 with molecular markers (IDH-wildtype/mutant, P53-wildtype/mutant, Ki67-high/low, and ATRX-retained/loss) was determined using the Mann-Whitney test. Kaplan-Meier survival analysis was applied to determine overall survival, and Cox regression analysis was performed to quantify the comparative hazard ratio. Patients were divided into groups (High and Low) using the median expression of miR-21 as the cut-off value. The log-rank test was also applied for survival analysis. The R statistical platform, base software version 3.3.2, and RStudio 2023.03.0+386 "Cherry Blossom" were used for statistical data analysis and graphical representation.

### Enrichment Analysis of mir-21 Target Genes and Pathways

First, we accessed the miRDB database which contains target genes of microRNAs and extracted a list of genes targeted by miR-21. To investigate how dysregulation of miR-21 might contribute to cellular functions, we used this list to conduct a functional enrichment analysis using the Metascape tool.We specifically focused on pathways related to cancer, cell cycle, and survival, employing KEGG pathways, WikiPathways and oncogenic signature gene sets. Enrichment analysis was performed to explore the biological processes associated with these gene groups, with an adjusted P-value cutoff of < 0.05 to ensure statistical significance.

## Results

### Clinical Characteristics of Glioma Patients

Our study group consisted of 90 glioma patients, with a male-to-female ratio of 64:26, indicating a predominance of male patients. The median age of the participants was 37 years. Based on the CNS5 WHO 2021 classification, the patients were categorized into glioma grades 1 to 4. Grade 4 included 48.8% of patients with histological types of glioblastoma, high-grade glioma IDH-wildtype and astrocytoma. Grade 3 included 12.2% of patients with astrocytoma and oligodendroglioma. Grade 2 included 26.6% of patients with astrocytoma, oligodendroglioma, and low-grade glioma with IDH wild type while 12.2% of patients were classified as grade 1 with histological types including pilocytic astrocytoma and ganglioglioma.

The molecular marker status showed that 56.6% of the patients had IDH1 wildtype, while 43.4% were IDH1 mutant. P53 mutations were observed in 53.5% of the cases, and 18.7% had lost ATRX expression. Notably, 59.3% of the tumors exhibited high Ki67 proliferation indices, indicating a high rate of cell division and potentially aggressive tumor behavior.

Regarding treatment, 62.2% of patients underwent chemotherapy and radiotherapy. A significant portion of the cohort (82.2%) had primary tumors at initial presentation, while 16.6% had recurrent tumors, and 1.1% experienced progression. The median tumor size was 35 cm³, indicating substantial variability in tumor burden among patients.

The pre-operative Karnofsky Performance Scale (KPS) scores indicated that 62.5% of patients had scores ≥80, reflecting relatively good functional status prior to surgery, while 7.5% had lower scores (<80). Post-operatively, 57.9 % of patients maintained KPS scores ≥80, whereas 42% had scores <80, suggesting that surgery had a variable impact on patient functionality.

Survival analysis showed that 58.4% of patients were alive at the time of the last follow-up, with a mortality rate of 41.5%. These survival rates underscore the critical need for effective therapeutic interventions, especially considering the high prevalence of aggressive glioma subtypes in the cohort. The detailed clinical parameters and characteristics of the glioma patients in our study are presented in Table 2.

**Table 2:** Clinicopathological characteristics of glioma patients in tissue samples.

| Clinicopathological features | No. (%) | Clinicopathological features | No. (%) |
| --- | --- | --- | --- |
| **Gender** |  | **Survival Status** |  |
| Male | 64 (71.11%) | Alive | 52 (58.4%) |
| Female | 26 (28.8%) | Deceased | 37 (41.5%) |
| **Glioma WHO Grades** |  | **Tumor Types** |  |
| Grade 1 | 11 (12.2%) | Pilocytic Astrocytoma | 8 (8.8%) |
| Grade 2 | 24 (26.6%) | Oligodendroglioma | 15 (16.6%) |
| Grade 3 | 11 (12.2%) | Glioblastoma | 31 (34.4%) |
| Grade 4 | 44 (48.8%) | High Grade Diffuse Glioma | 4 (4.4%) |
| Normal | 10 | Ganglioglioma | 2 (2.2%) |
| **Age (years) Median (range)** | 37(years) | Low Grade Diffuse Glioma | 4 (4.4%) |
| **Molecular Marker status** |  | Astrocytoma | 26 (28.8%) |
| IDH1 Mutant | 36 (43.4%) | **Radiotherapy** |  |
| IDH1 Wildtype | 47 (56.6%) | Yes | 56 (62.2%) |
| P53 Wildtype | 39 (46.4%) | No | 29 (32.2%) |
| P53 Mutant | 45 (53.5%) | NA | 5 (5.5%) |
| ATRX Retained | 65 (81.2%) | **Chemotherapy** |  |
| ATRX lost | 15 (18.7%) | Yes | 56 (62.2%) |
| Ki67 High | 51 (59.3%) | No | 29 (32.2%) |
| Ki67 Low | 35 (40.6%) | NA | 5 (5.5%) |
| **Pre-operative KPS score** |  | **Recurrence** |  |
| >=80 | 55 (62.5%) | Primary | 74 (82.2%) |
| <80 | 33 (7.5%) | Recurrence | 15 (16.6%) |
| **Post-operative KPS score** |  | Progression | 1 (1.1%) |
| >=80 | 51 (57.9%) | **Average Tumor Size** |  |
| <80 | 37 (42%) | 35 cm<sup>3</sup> |  |

Table 2 provides a detailed breakdown of these clinicopathological features, highlighting significant factors that may influence patient prognosis and treatment outcomes. The presence of key molecular markers like IDH1, P53, and ATRX, as well as high Ki67 indices, suggests avenues for personalized treatment approaches that could potentially improve survival and quality of life for glioma patients.

### miR-21 Gene Expression in Age and Gender Subgroups

To understand the dynamics of tumor-associated miR-21 expression in tissue samples, we conducted a comparative analysis based on age and gender. Patients were categorized into two age groups: those below 50 years and those 50 years or older. Our results demonstrated significantly elevated miR-21 expression in patients aged 50 years and above (p<0.001) suggesting a potential age-related increase in miR-21 levels (Fig. 1A). However, since the distribution of glioma grades was not uniform between the two age groups, this finding may partly reflect the higher average glioma grade in patients over 50 years. In contrast, there was no significant difference in miR-21 expression between male and female patients, indicating a similar expression pattern across genders (Fig. 1B).

**Figure 1a:**
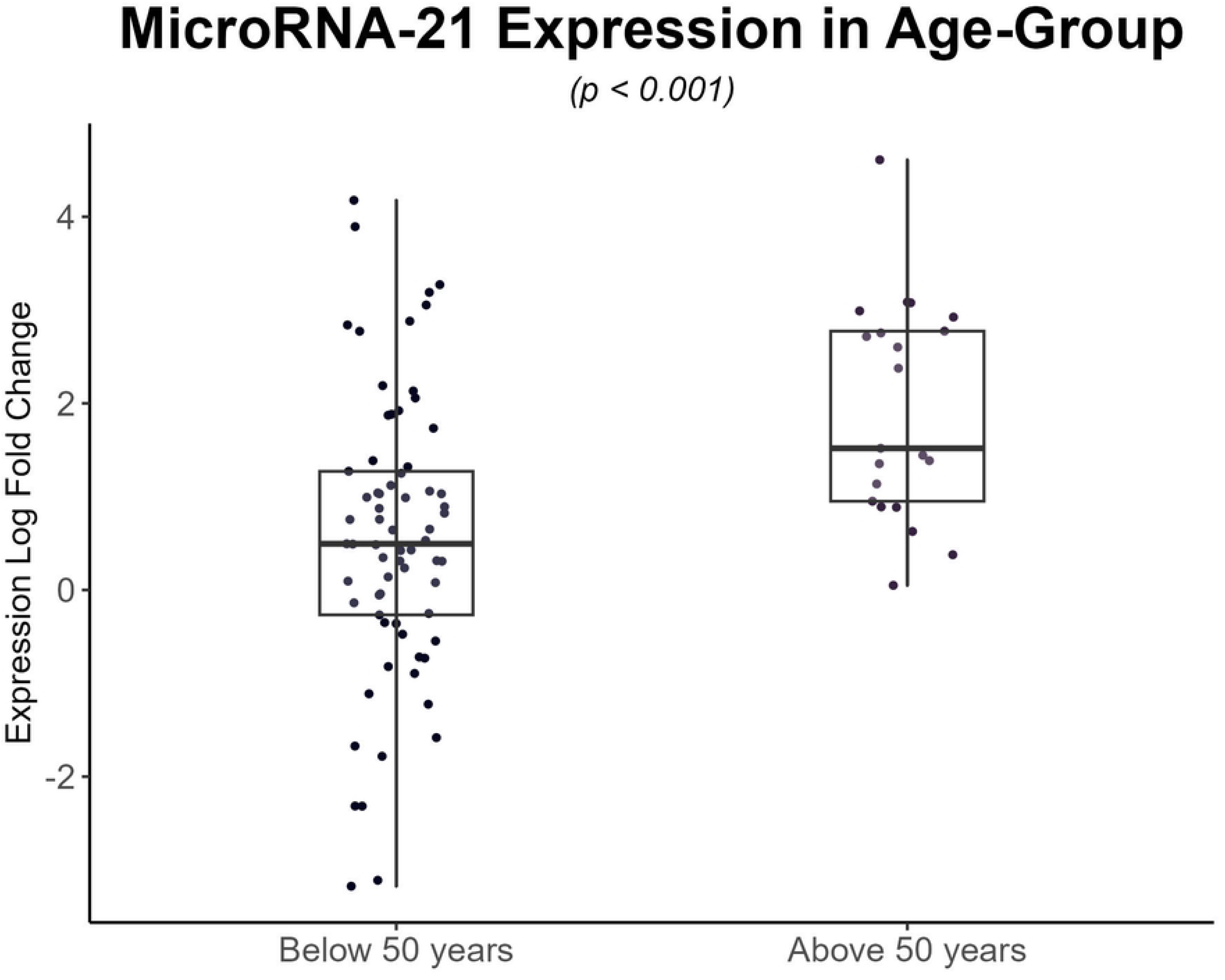
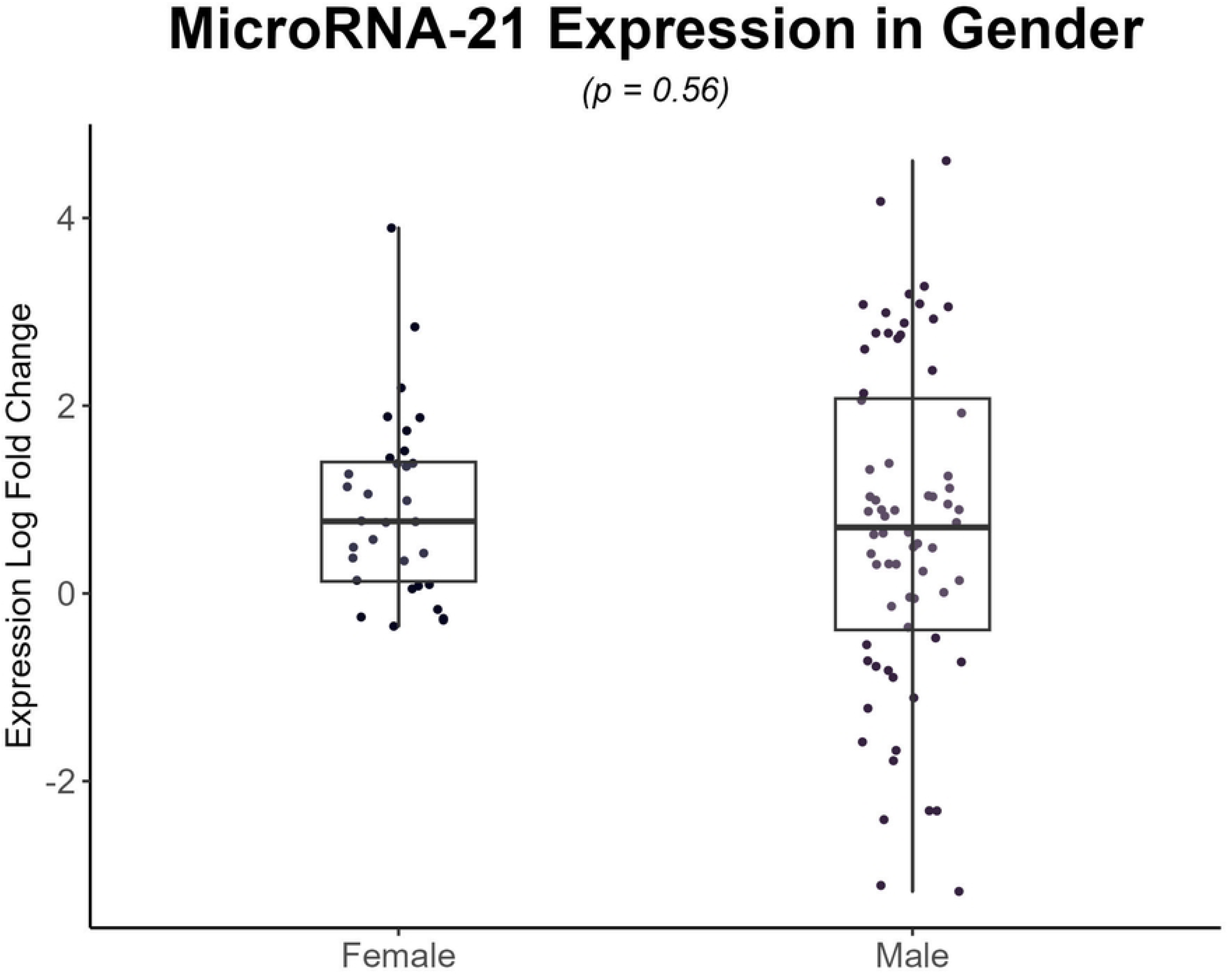
Illustrates age-dependent variation in miR-21 expression levels among glioma patients, comparing those aged 50 years and above with those below 50 years. Elevated miR-21 expression was notably observed in patients aged 50 years and above (p<0.001). Figure 1b: presents gender-dependent analysis of miR-21 expression, showing comparable levels between male and female patients, indicating no significant difference in miR-21 expression based on gender.

### miR-21 Expression in Tumor Regions and Volume of Glioma Patients

In our study, the tissue expression of miR-21 varied across different tumor regions. High expression levels were observed in the Bi-Frontal, right intraventricular region, followed by the right parietal and right temporal regions. Conversely, miR-21 expression was down-regulated in tumors located within the supra sellar and left temporal parietal regions (Fig. 2a). To further explore the relationship between miR-21 expression and tumor characteristics, we conducted a correlation analysis with tumor volume using Pearson’s correlation test. Our findings revealed a pattern of slight positive correlation between miR-21 expression levels and tumor volume, although this correlation was not statistically significant (r=0.15, p=0.163) (Fig. 2b).

**Figure 2:**
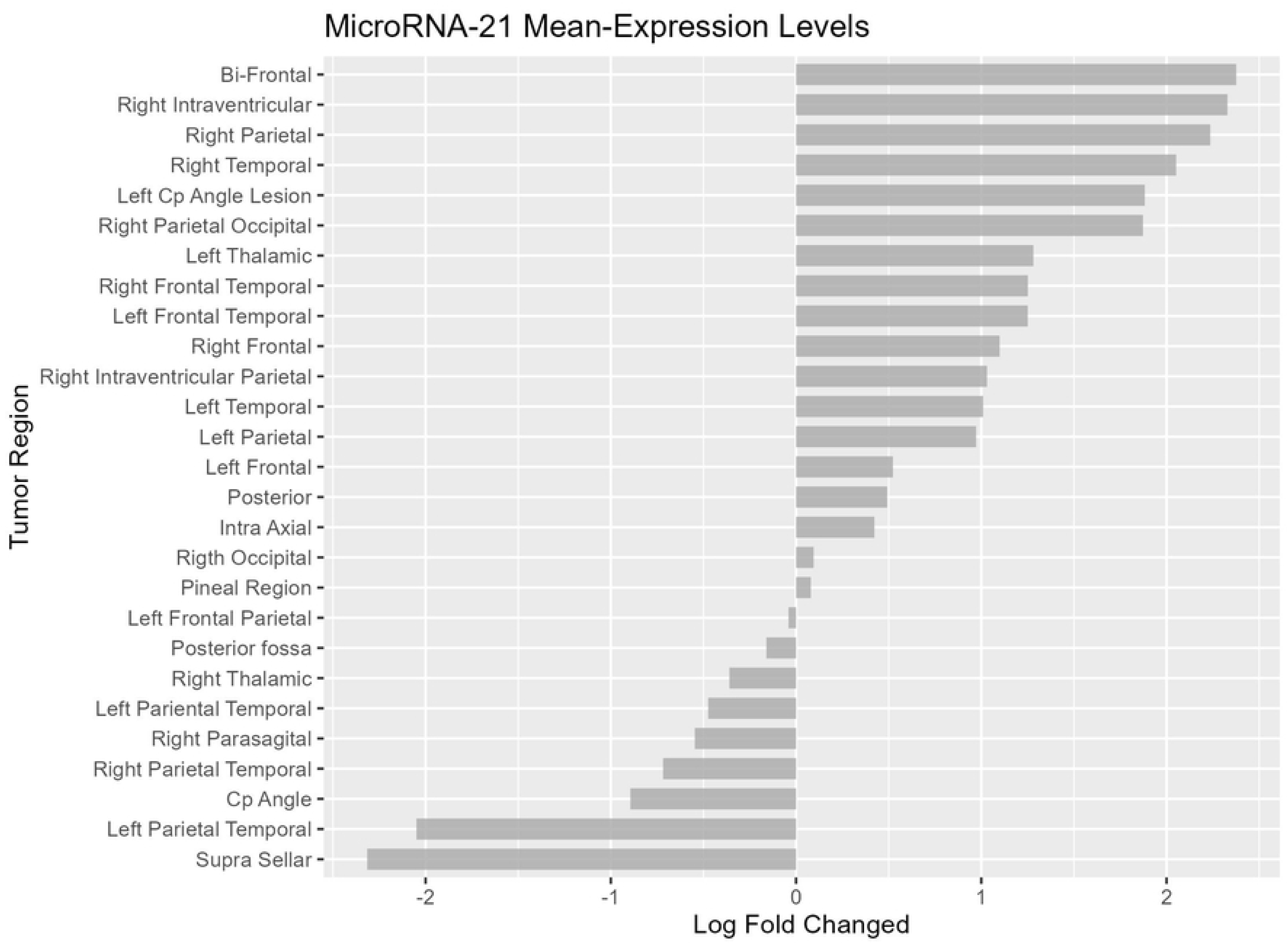
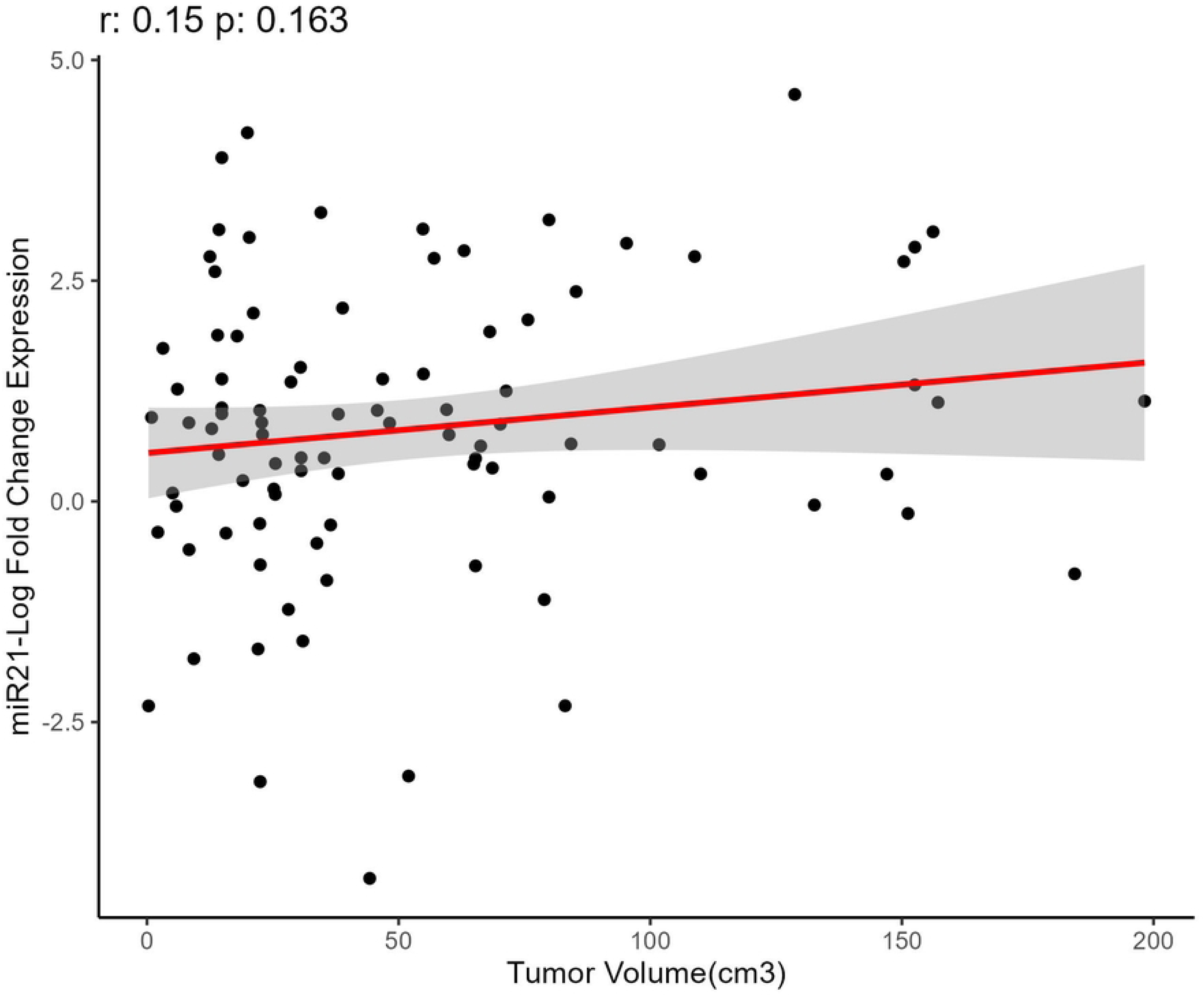
miR-21 expression analysis across glioma tumor region and volume. Fig. 2a: Depicts the mean miR-21 expression levels across different brain regions affected by glioma. Fig. 2b: Illustrates the correlation analysis between miR-21 expression levels and tumor volume (cm³).

**Figure 3:**
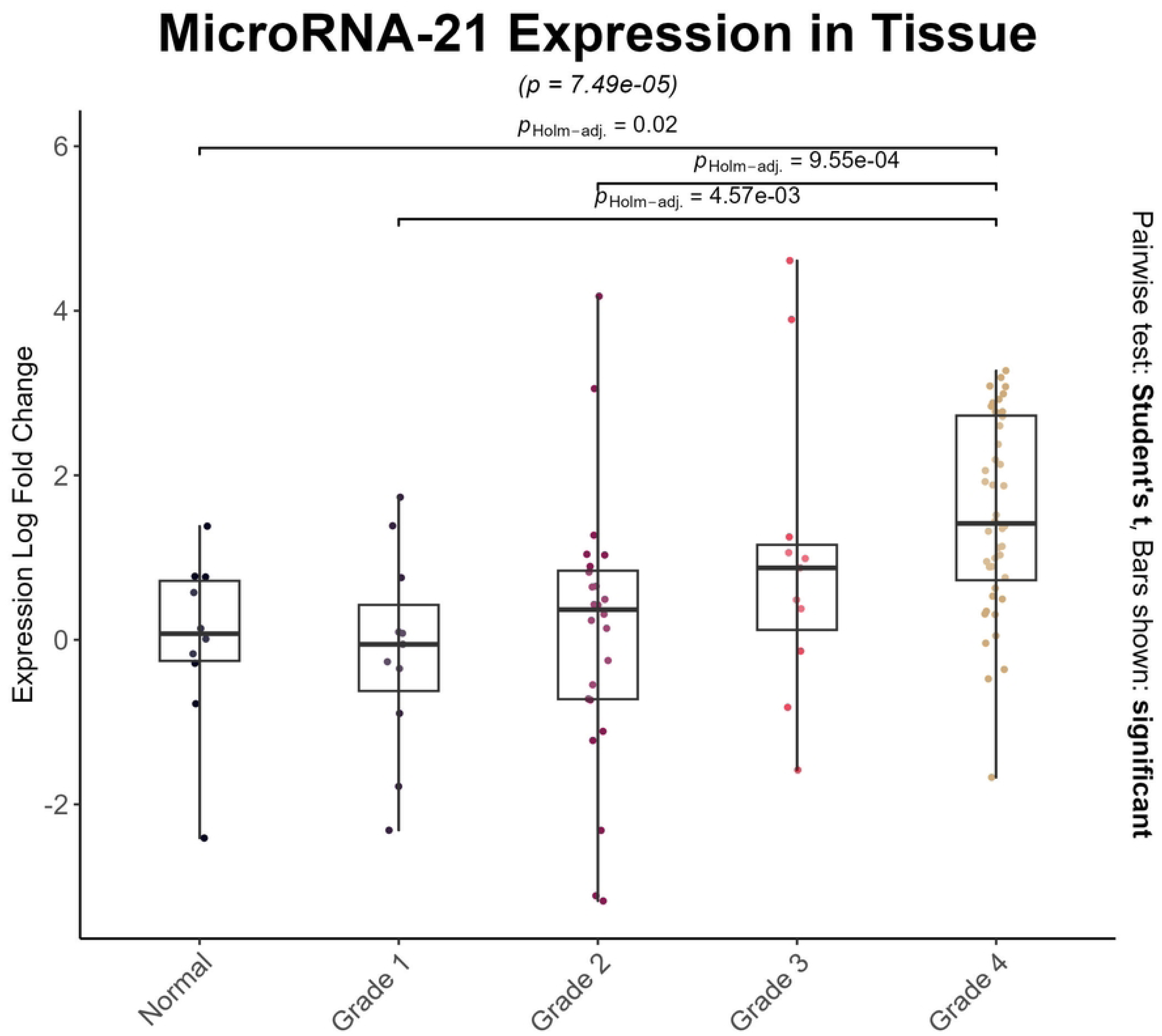
Illustrates the comparison of miR-21 expression levels in tissue samples from glioma patients across different grades, as well as normal tissues.

### mir-21 Tissue Expression Profile Across Different Grades of Glioma

Several studies have established that miR-21 expression is elevated in higher glioma grades. To investigate this trend, we compared miR-21 expression across various glioma grades and control tissues. For this analysis, we utilized a multivariate ANOVA along with pairwise comparisons (Holm-adjusted p-values) to evaluate differences between control tissues and glioma grades. Our findings demonstrated that miR-21 expression was significantly upregulated in grade 4 gliomas compared to control tissues (p = 0.02), grade 1 gliomas (p = 4.57e-03), and grade 2 gliomas (p = 9.55e-04), highlighting a clear association between increased miR-21 expression and higher glioma grades.

### Association of Molecular Markers with Mir-21 Expression in Glioma Patients

The latest classification system for gliomas leverages established molecular markers that are routinely tested for diagnostic purposes. We examined the expression difference of miR-21 with key molecular markers, including ATRX (loss/retained), IDH (wildtype/mutant), Ki-67 (low/high), and p53 (wildtype/mutant). Our analysis revealed a significant differential expression of miR-21 in patients with IDH-wildtype gliomas (p=0.03) and those with high Ki-67 expression levels (p < 0.001). In contrast, no significant differences in miR-21 expression were found in relation to ATRX status (p=0.89) or p53 status (p=0.90) (Fig. 4).

**Figure 4:**
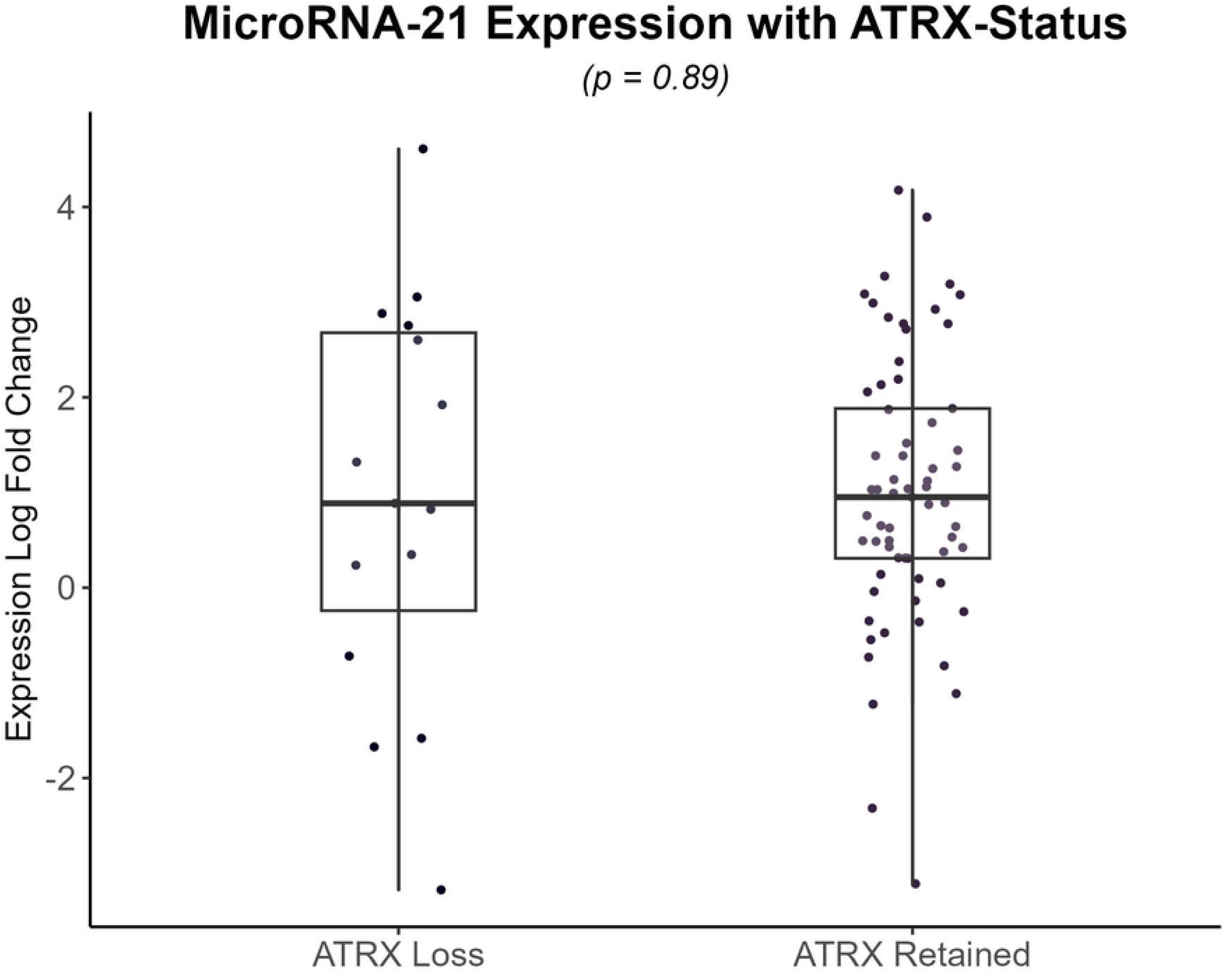
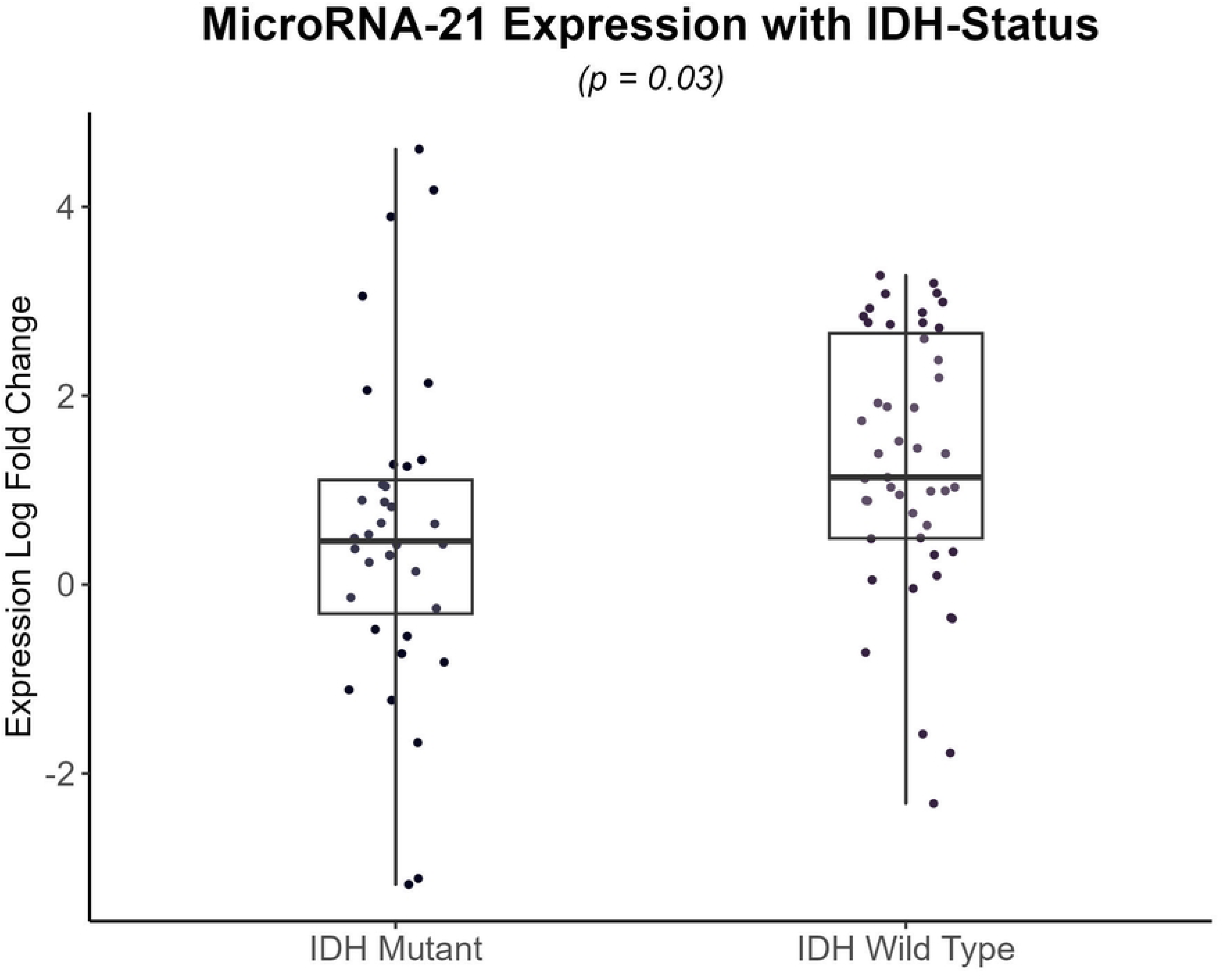
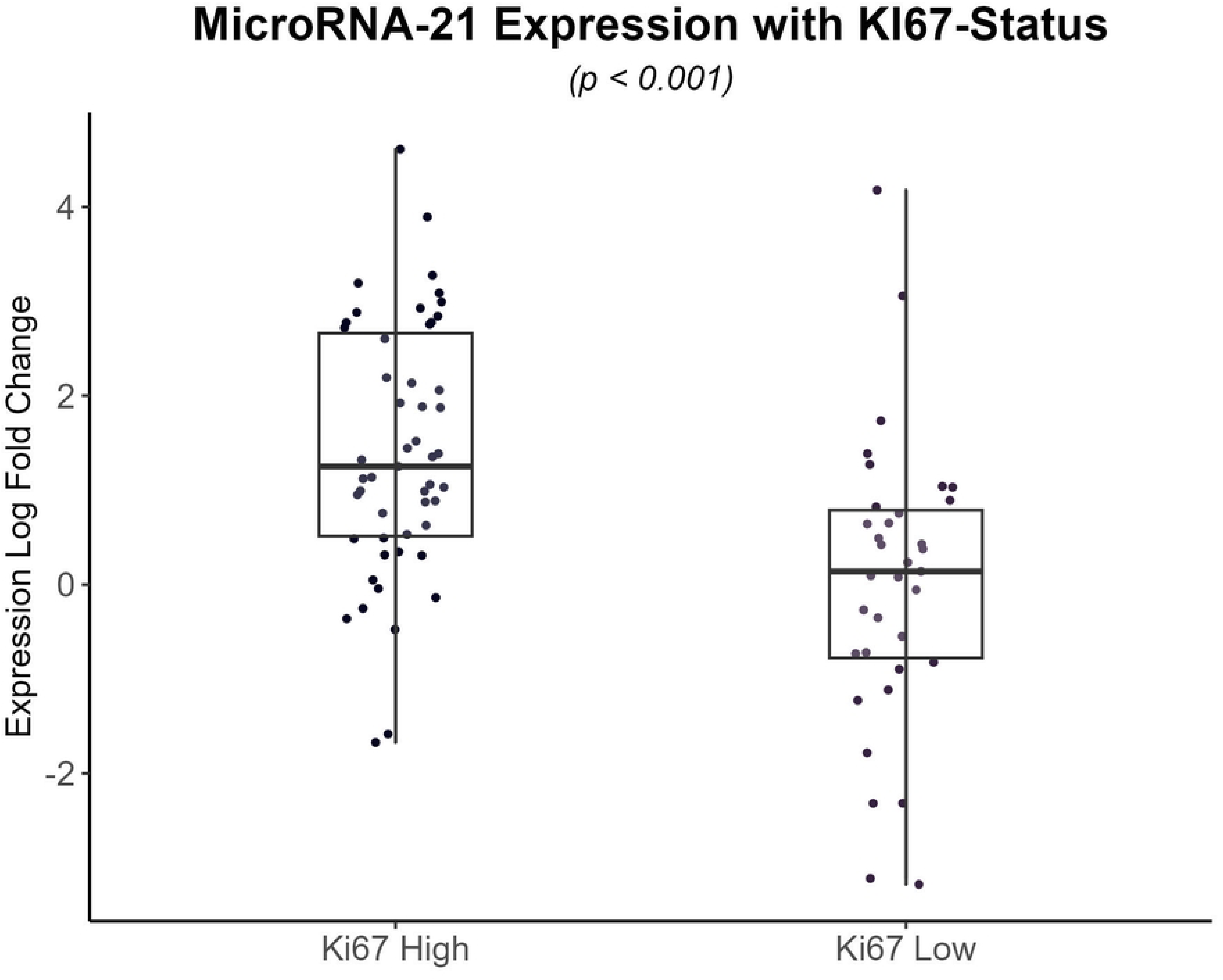
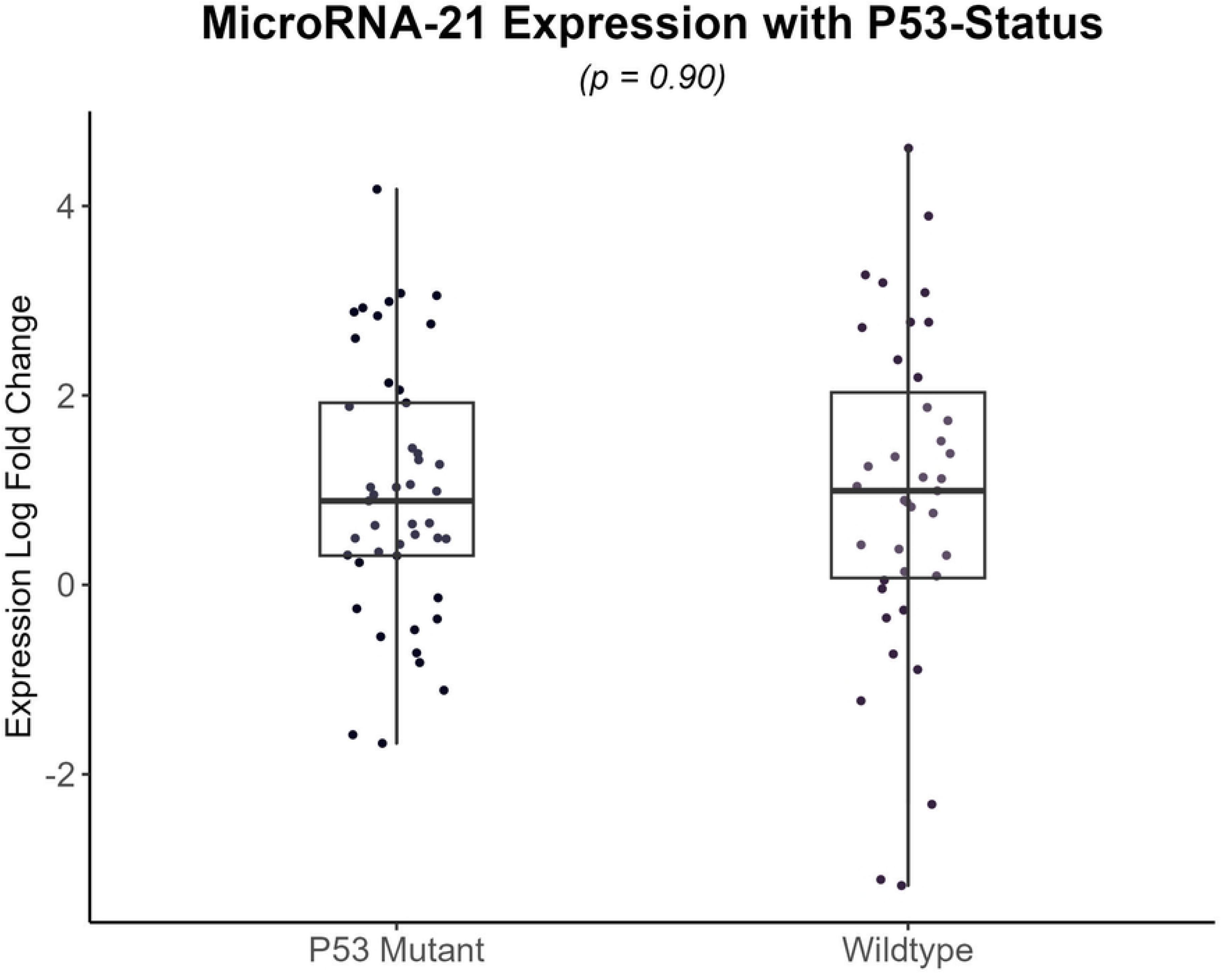
Expression analysis of miR-21 in relation to molecular marker status: ATRX (Loss and Retained), IDH (Wildtype and Mutant), Ki-67 (High and Low), and p53 (Wildtype and Mutant)

### Association of mir-21 with Overall Survival, Hazard Ratio and ROC Curve for Expression Prediction in Glioma Patients

To determine whether the expression of miR-21 could be used as prognostic biomarker, the patients were divided into low and high expression groups according to miR-21 median expression. Within 5 years of follow-up period, the mortality rate for the cohort was 41.5 % (37 patients). Glioma patients from the low miR-21 expression group showed significantly higher overall survival (log-rank = 0.00046) (Fig 5a). Moreover, a quantitative evaluation of the hazard ratio was done using the Cox regression method in miR-21 high and low expression groups.. The analysis indicated that patients from the high expression group were at 3.4 times higher risk of mortality (95% confidence interval: upper 7.1, lower 1.6), in comparison to patients in the low expression group (HR: 3.4, log rank p < 0.001) (Fig 5b). ROC curves for miR-21 were generated with mortality during the follow-up period as the primary endpoint, demonstrating its potential as a valuable prognostic biomarker (AUC = 0.74) (Fig 5c). This matches the prognostic accuracy of currently used molecular markers (IDH, P53, and ATRX), suggesting that integrating miR-21 expression status with existing testing methods could offer enhanced predictive insights into disease progression, helping to inform treatment decisions.

**Figure 5:**
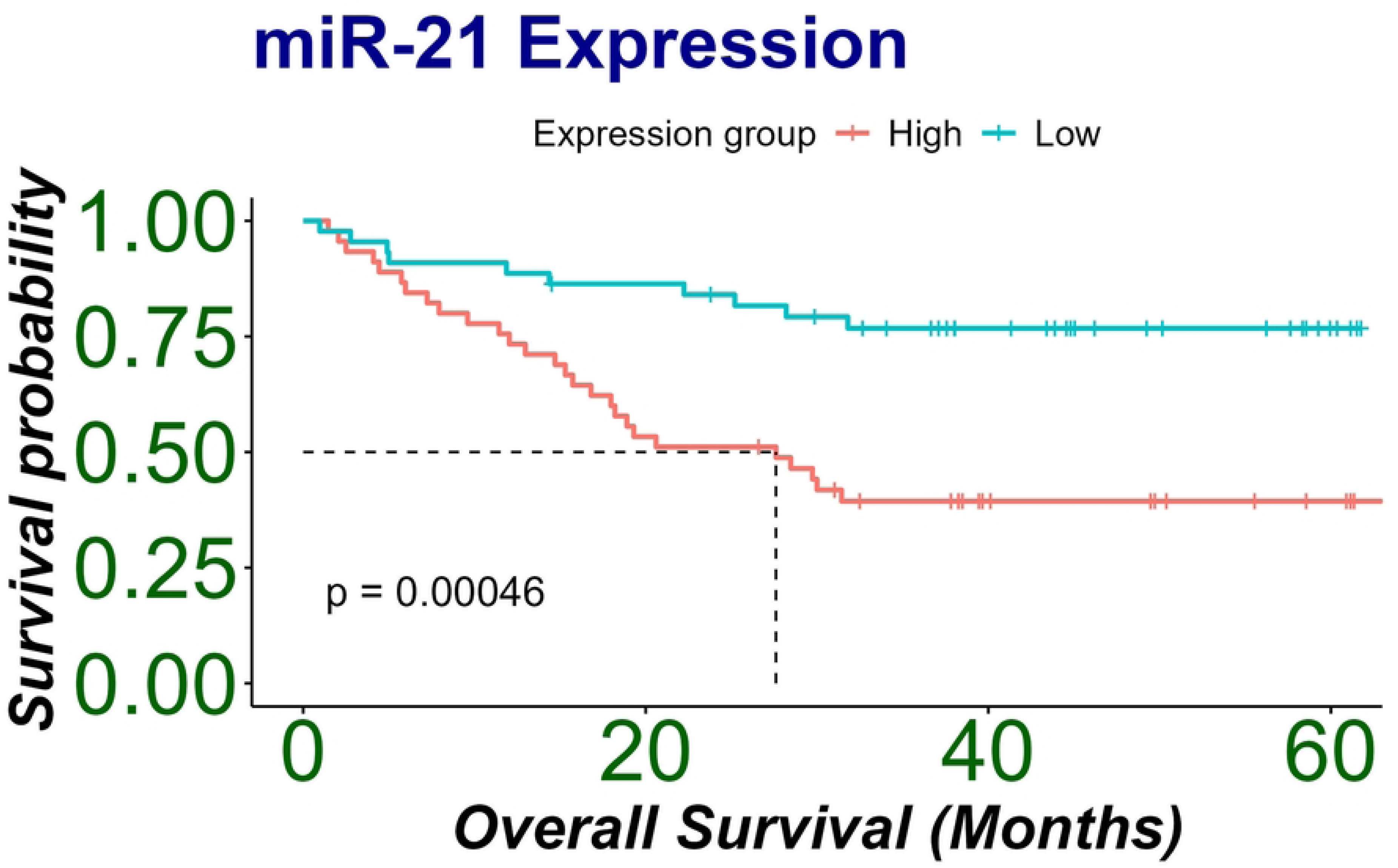
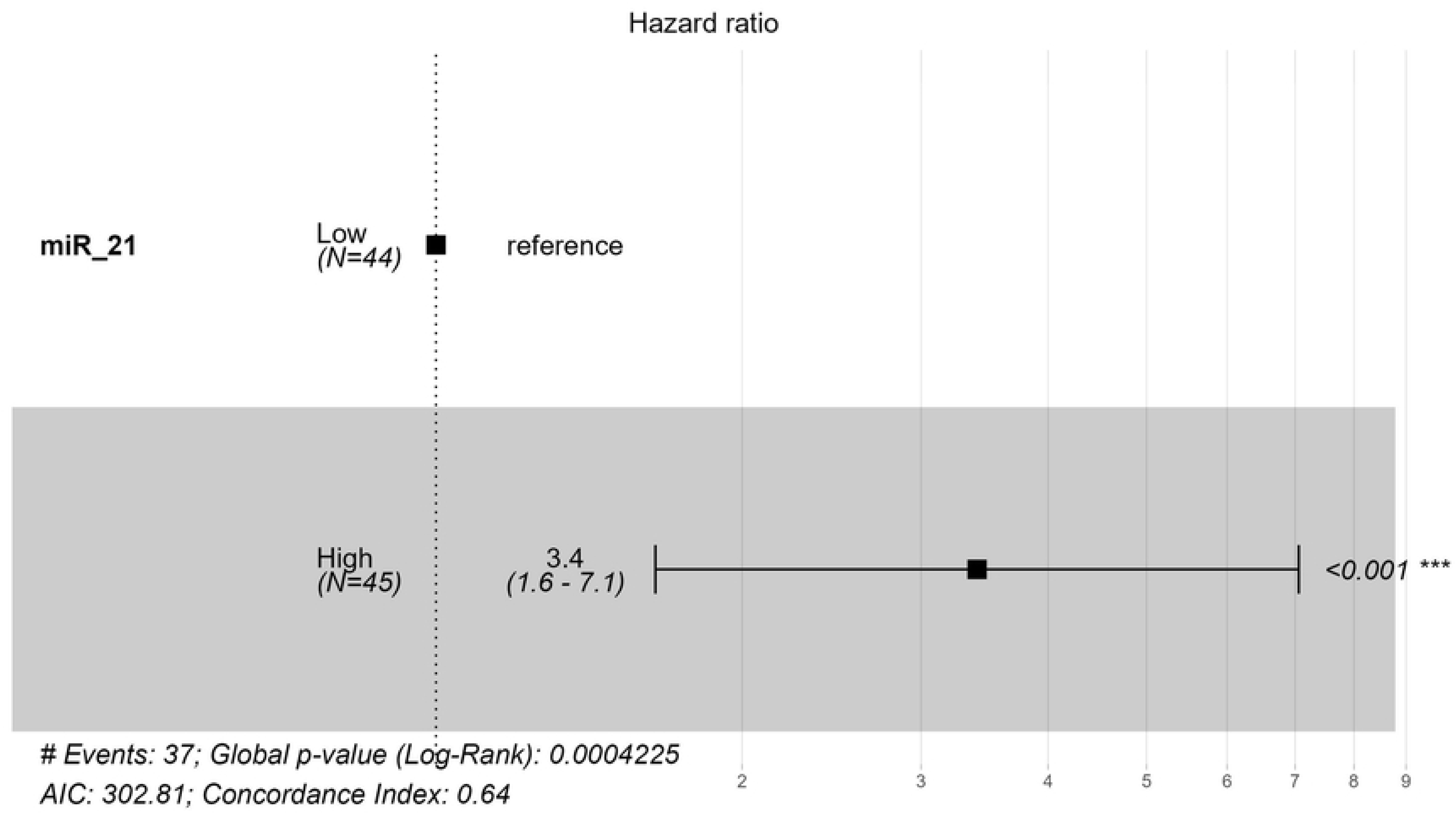
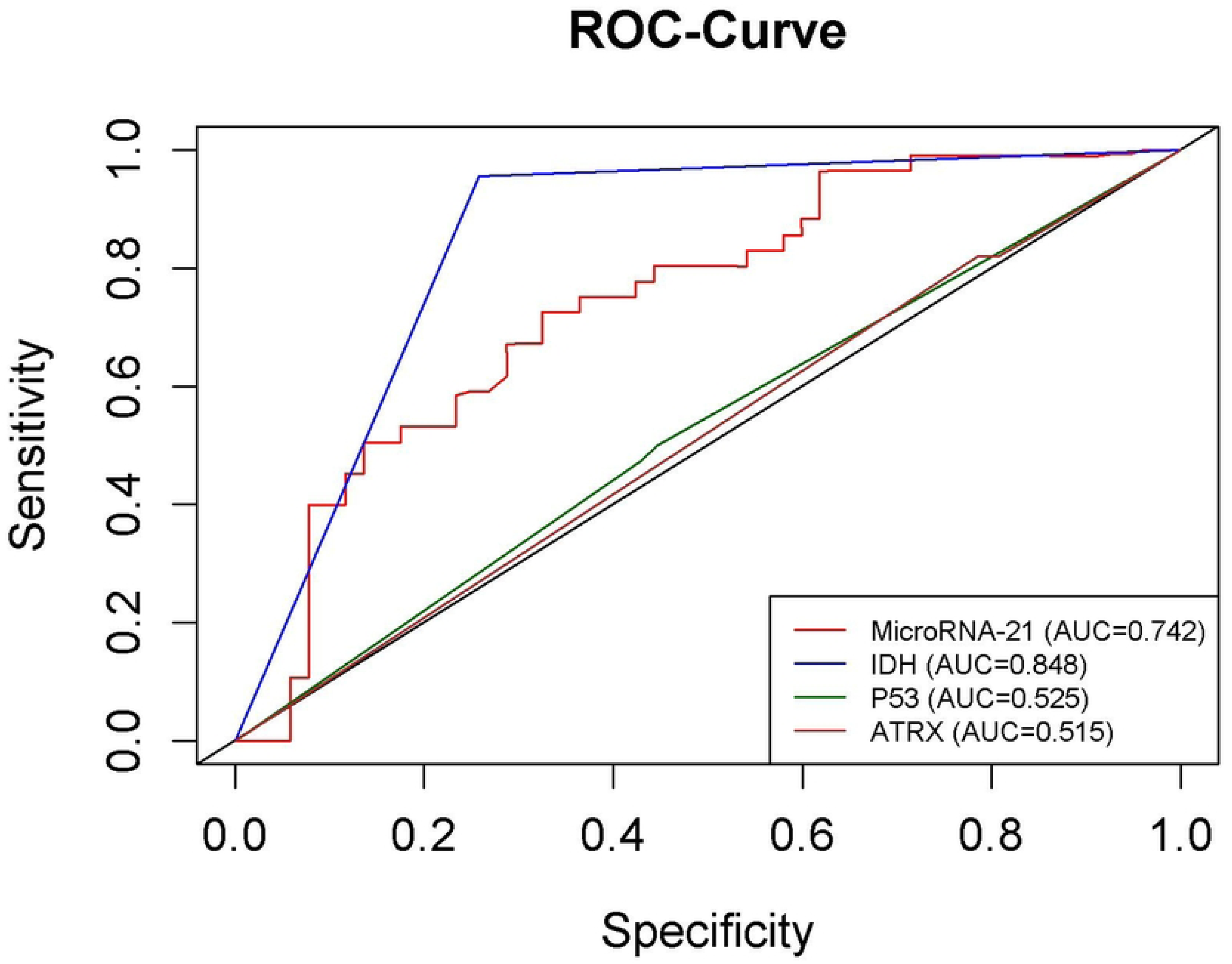
Kaplan-Meier survival curves of glioma patients in miR-21 high and low expression groups (Fig.5a). Hazard ratio of glioma patients in miR-21 high and low expression group (Fig.5b). Receiver operating characteristic (ROC) curves indicating prognostic performance of miR-21 expression and existing molecular markers (IDH, P53 and ATRX) (Fig.5c).

### Overall Survival Pattern in Glioma Patient with Molecular Markers and Mir-21 Expression

To understand the combined influence of miR-21 expression levels and molecular marker status on overall survival (OS) of the patients, we compared miR-21 expression groups and molecular marker mutations, in various combinations as shown in Fig 6a, high miR-21/IDH-mutant, low miR-21/IDH-mutant, high miR-21/IDH-wildtype, and low miR-21/IDH-wildtype; (Fig 6b) high miR-21/Ki-67 high, low miR-21/Ki-67 high, high miR-21/Ki-67 low, and low miR-21/Ki-67 low; (Fig 6c) high miR-21/p53-mutant, Low miR-21/p53-mutant, high miR-21/p53-wildtype, and low miR-21/p53-wildtype and (Fig 6d) high miR-21/ATRX loss, low miR-21/ATRX loss, high miR-21/ATRX retained, and low miR-21/ATRX retained.

**Fig 6:**
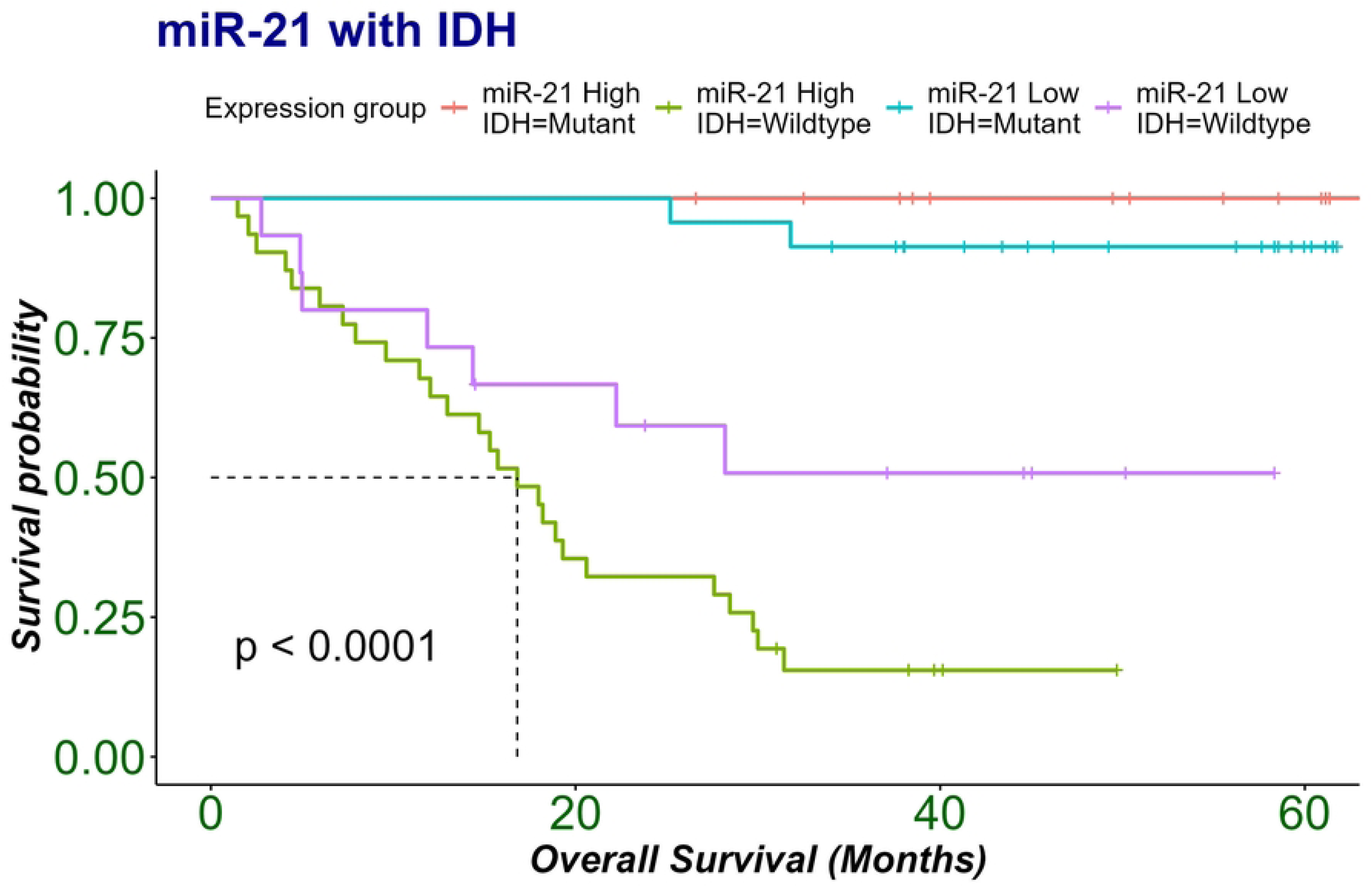
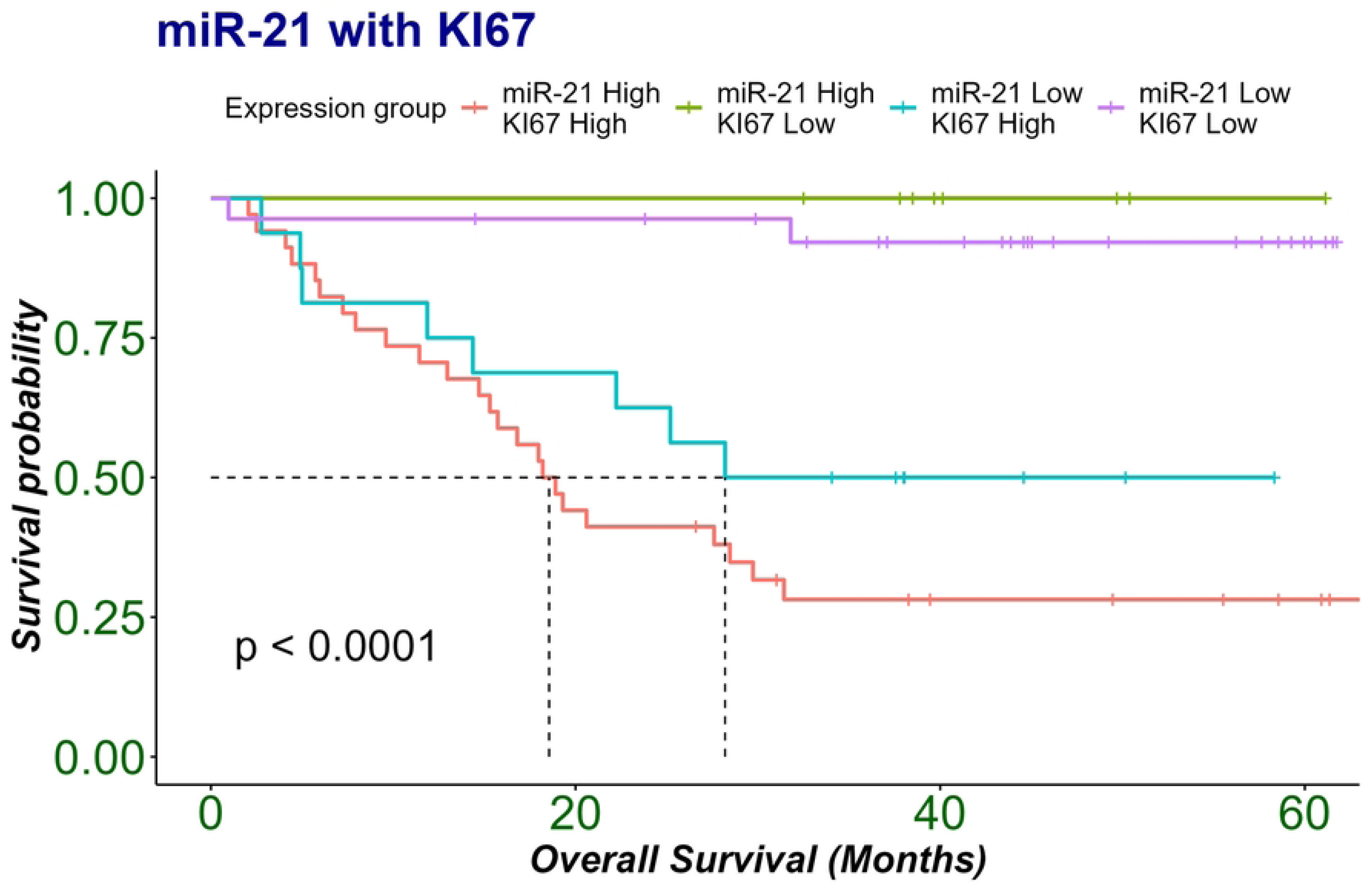
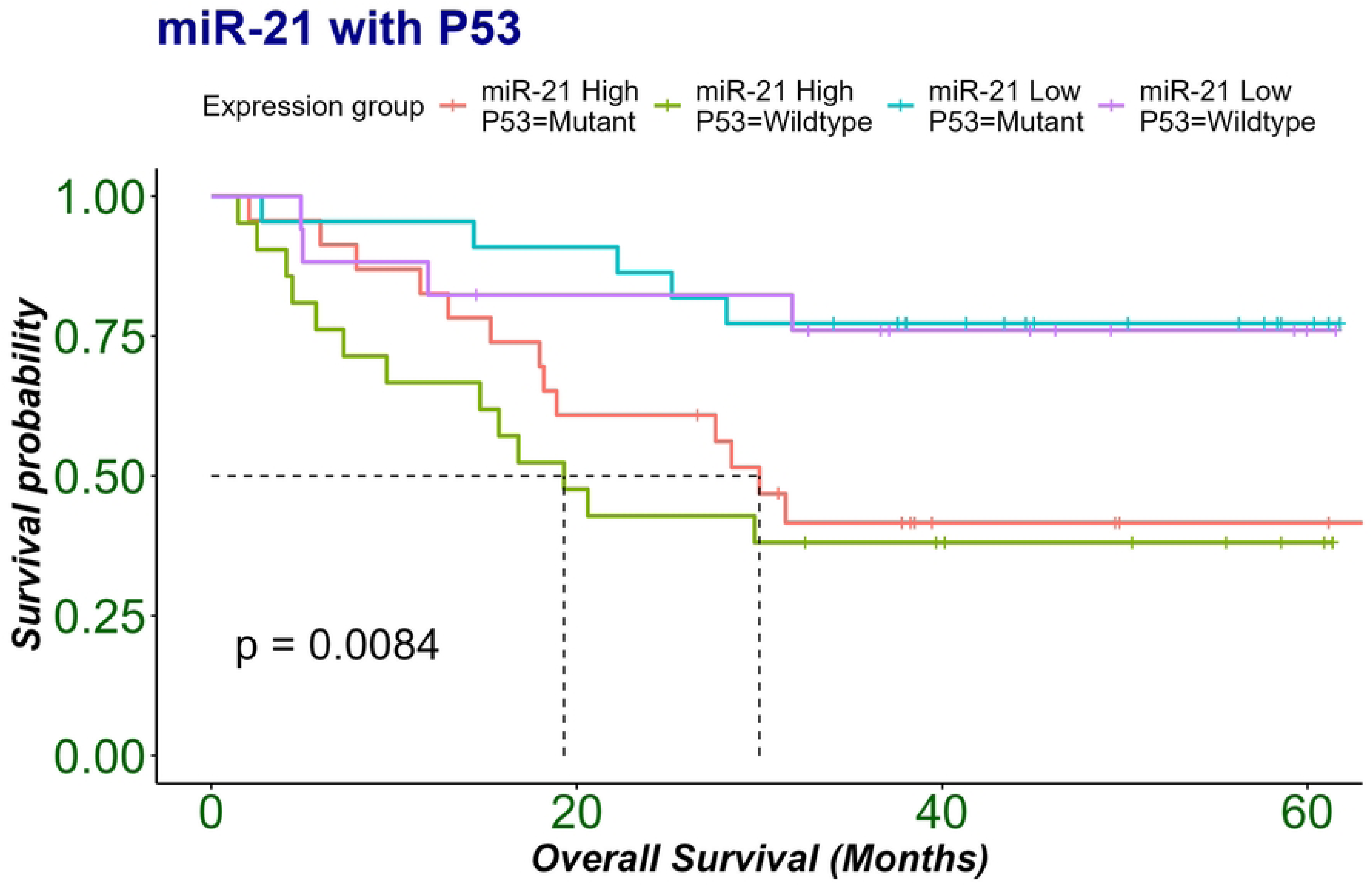
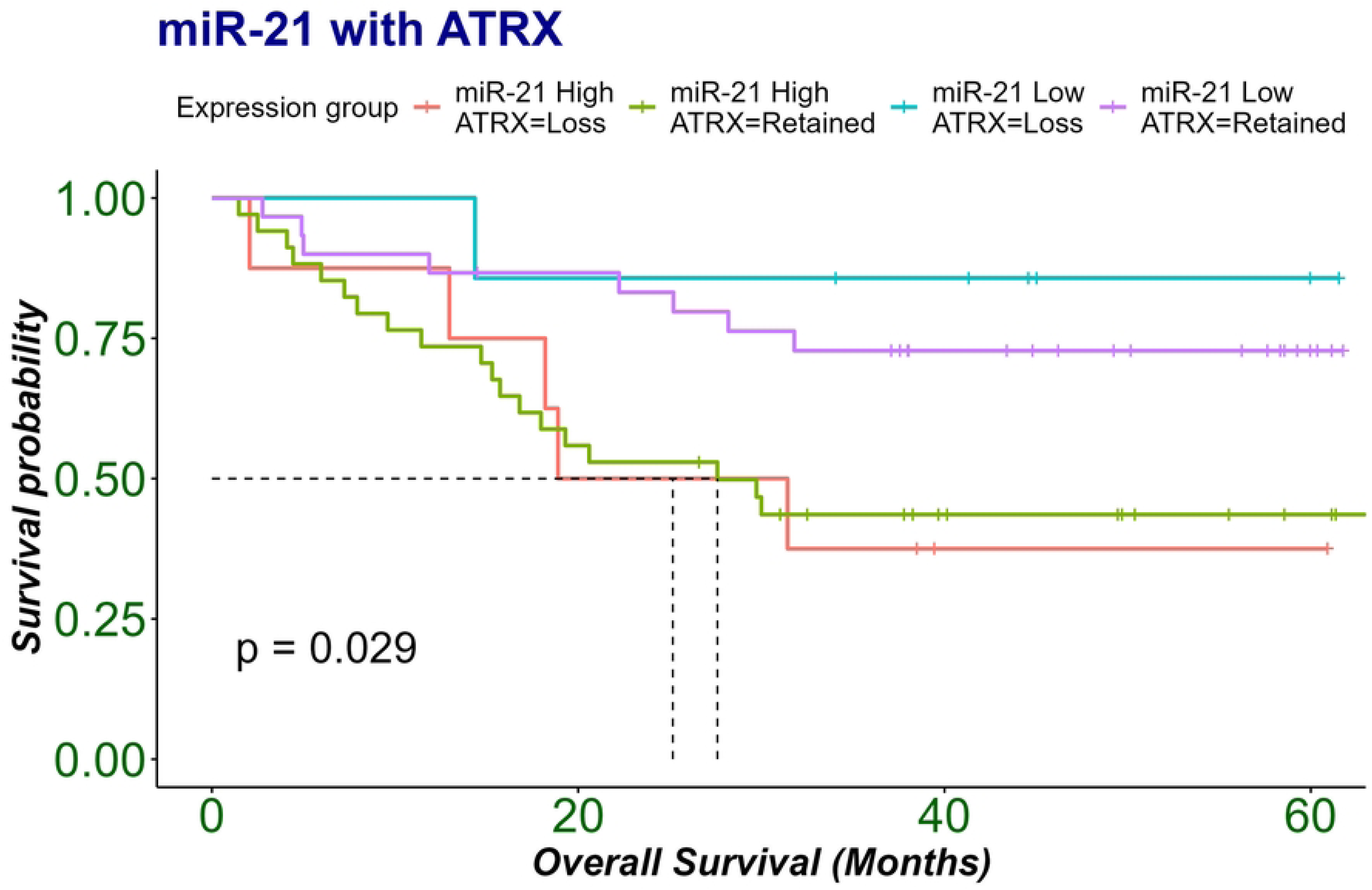
Kaplan-Meier survival curves for combined group miR-21 expression and molecular markers, 5-year overall survival. (Fig6a) IDH (wild-type and mutant) with high and Low miR-21 expression group (Fig 6b) Ki-67 (high and low) with high and Low miR-21 expression group (Fig 6c) p53 (wildtype and mutant) with high and Low miR-21 expression group (Fig 6d) ATRX (lost and retained) with High and Low miR-21 expression group

The results showed a strong association between high-miR-21 expression group and IDH-wildtype patients (p<0.0001), high-Ki67 patients (p<0.0001), p53 wildtype patients (p<0.0084) and ATRX retained patients (p=0.029) with poor overall survival. Thus, indicating that, high miR-21 expression is an independent prognostic indicator of poor overall survival when comparing with molecular markers.

For comparative analysis of miR-21 expression in tumor tissues and peripheral samples, the expression profiles in serum samples were also analyzed. For this we collected serum samples from glioma patients, these are overlapping serum samples from the patients we collected tissue samples as well as some new recruits for the study. The discordance between the number of tissue and serums samples can be attributed to the fact that some samples could not be timely collected, while some patients were lost to follow-up postoperatively. Moreover, few samples were not included due to insufficient serum aliquoting from blood samples. Serum samples were collected from a total of 42 patients pre and post operatively while samples from 8 healthy individuals were also collected as controls. The clinical parameters of this group are listed in table 3.

**Table 3:** Clinicopathological characteristics of glioma patients in serum samples.

| **Clinicopathological features** | **No. (%)** | **Clinicopathological features** | **No. (%)** |
| --- | --- | --- | --- |
| **Gender** |  | **2021 WHO CNS5 grades** |  |
| Male | 27 (64.2%) | Grade 1 | 6 (14.28%) |
| Female | 15 (35.7%) | Grade 2 | 5 (11.90%) |
| **Age (years)** |  | Grade 3 | 5 (11.90%) |
| Medians (range) in months | 37.5 | Grade 4 | 26 (61.90%) |
| **Survival Status at 4 years** |  | Normal | 8 |
| Alive | 21 (50%) | **Molecular Marker Status** |  |
| Dead | 21 (50%) | IDH1 Mutant | 14 (36.8%) |
| **Histological type (WHO CNS 2021)** |  | IDH1 Wildtype | 24 (63.1%) |
| Pilocytic Astrocytoma | 4 (9.5%) | ATRX Retained | 24 (88.88%) |
| Oligodendroglioma | 4 (9.5%) | ATRX Lost | 12 (33.33%) |
| Glioblastoma | 17 (40.4%) | p53 Mutant | 21 (55.2%) |
| Ganglioglioma | 2 (4.7%) | p53 Wildtype | 17 (44.7%) |
| Astrocytoma | 11 (26.1%) | Ki-67 High | 30 (73.17%) |
| High Grade Glioma | 4 (9.5%) | Ki-67 Low | 11 (26.82%) |
| Tumor Volume (Median cm3) |  |  |  |

### miR-21 Serum Expression Profile Across Different Glioma Grades

RT-qPCR was employed to analyze the miR-21 expression profiles in serum samples of glioma patients in pre- and post-surgery serum samples. They were also assessed for variations using Multivariant ANOVA along with pairwise test (Holmes adjusted p value) for intercomparison of miR-21 expression in control and across different grades of glioma. The same trend as in tissue samples was observed in preoperative serum samples, a statistically significant difference was found in pre-op grade-2 (p= 0.0031), pre-op grade-3 (p= 0.0031), and pre-op grade-4 (p= 0.033) when compared with control. Furthermore, comparison between pre-op and post-op serum revealed significant decrease in miR-21 expression after surgery in grade 2 (p= 0.0079), grade 3 (p= 0.0079) and grade 4 (p= 0.0096) glioma patients (Fig 7).

**Figure 7:**
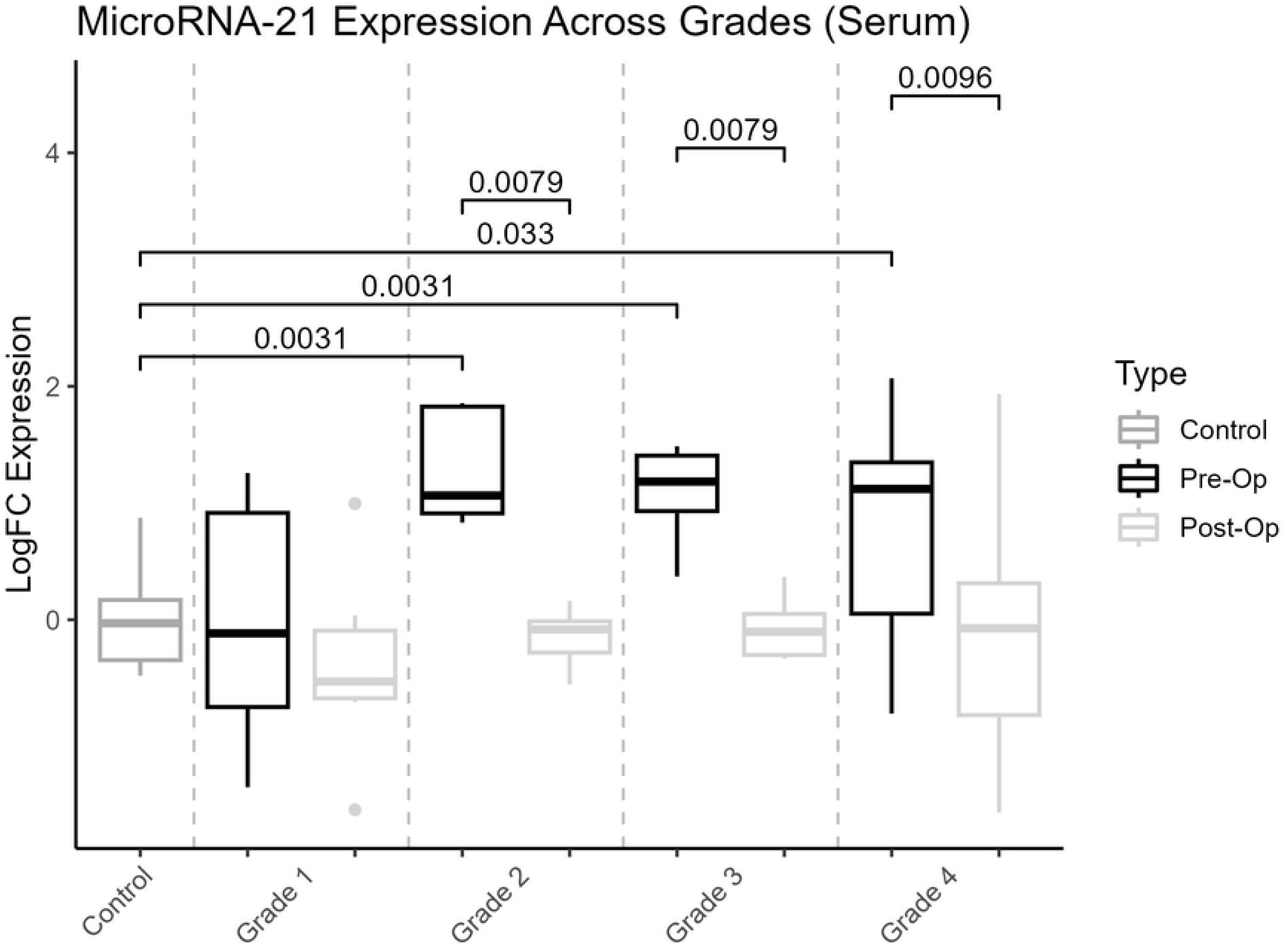
Trend in the miR-21 expression levels in serum samples from glioma patients and control.

### miR-21 expression pattern in glioma patient’s serum based on tumor region and tumor volume

Similarly, Mean expression value of miR-21 according to tumor region are represented in Figure 8A and miR-21 expression shows positive correlation with tumor volume, although this correlation was not significant (r=0.37, p=0.139) in Figure 8B.

**Figure 8:**
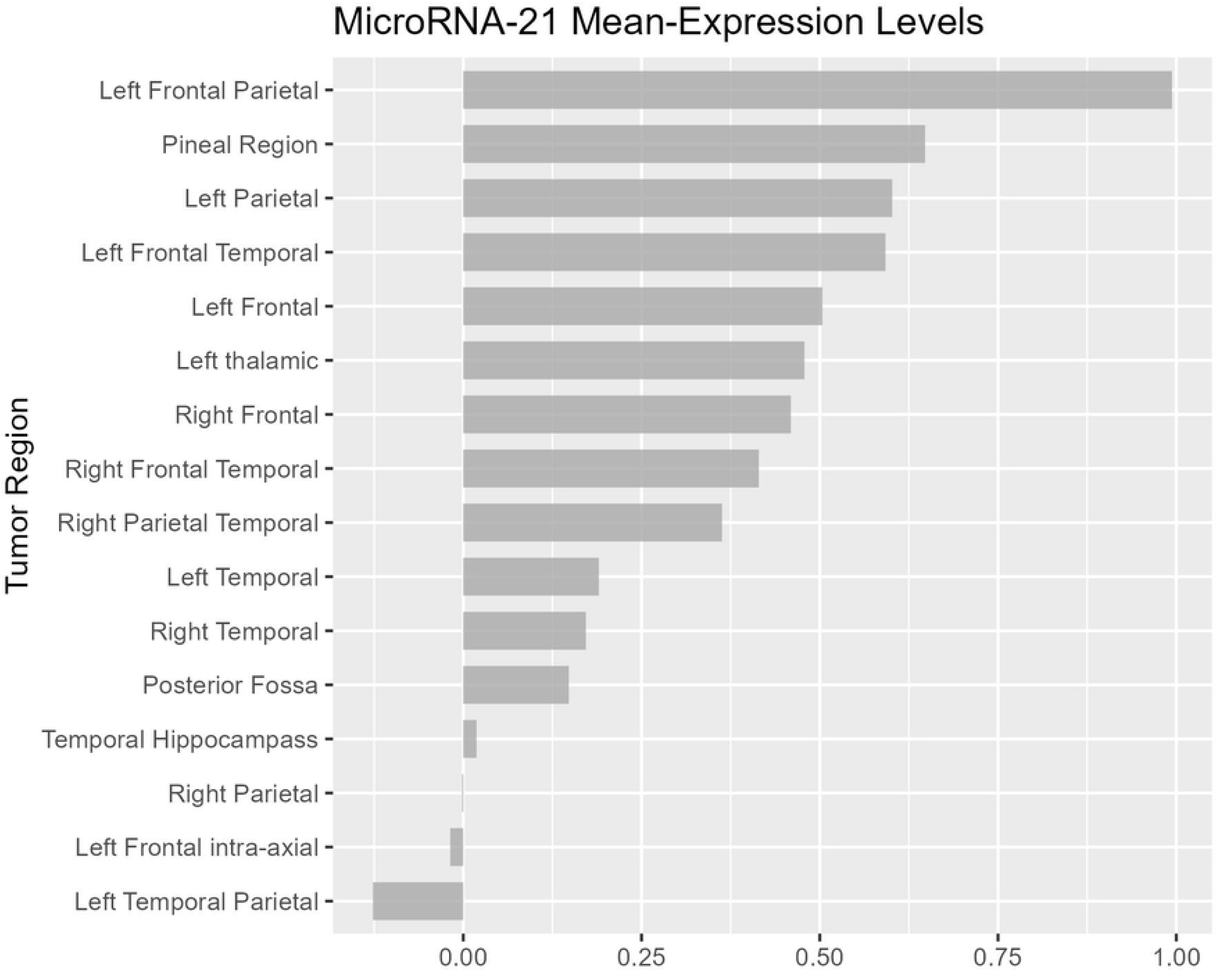
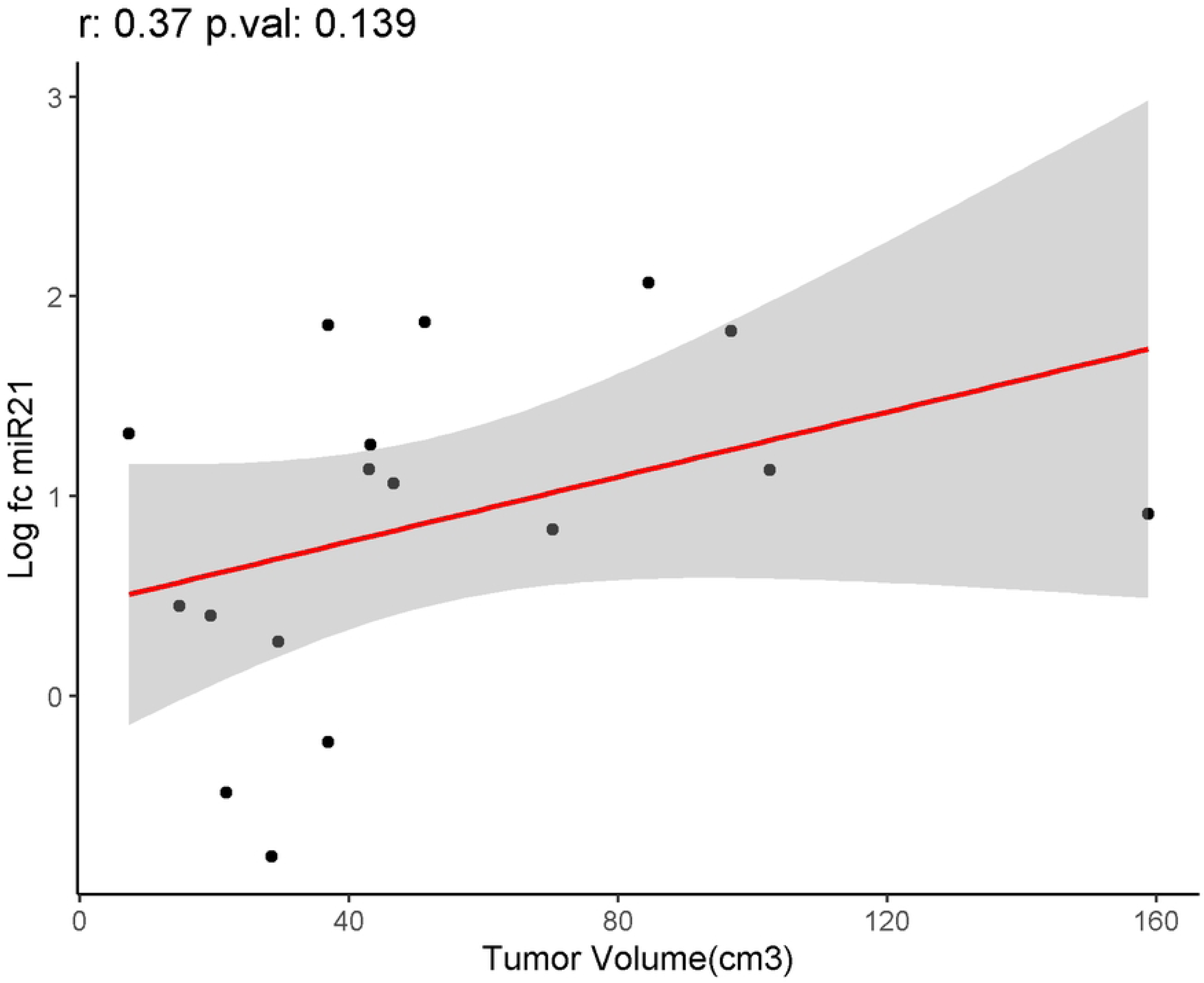
miR-21 expression analysis across glioma tumor region and volume (Fig.8a) miR-21 mean expression level of glioma in different brain regions. (Fig.8b) Correlation analysis of miR-21 serum expression level and tumor volume (cm^3^)

### Association of Molecular Markers with miR-21 Serum expression in Glioma Patients

Figure 9 examines associations of molecular markers with serum miR-21 expression, pre- and post-operative glioma patients. ATRX-retention in gliomas is significantly associated with lower miR-21 expression (p=0.0003) in serum after surgery, within our cohort. IDH-mutant (p=5e-08) and –wildtype (p=0.039) gliomas show significant decrease in miR-21 expression after surgery in patients with mutant as decrease pattern noted in IDH-wildtype same pattern were followed in patients with P53 mutant (p=0.00082) and wild type (p=0.08508), along with high Ki-67 (p=0.0026) gliomas.

**Figure 9:**
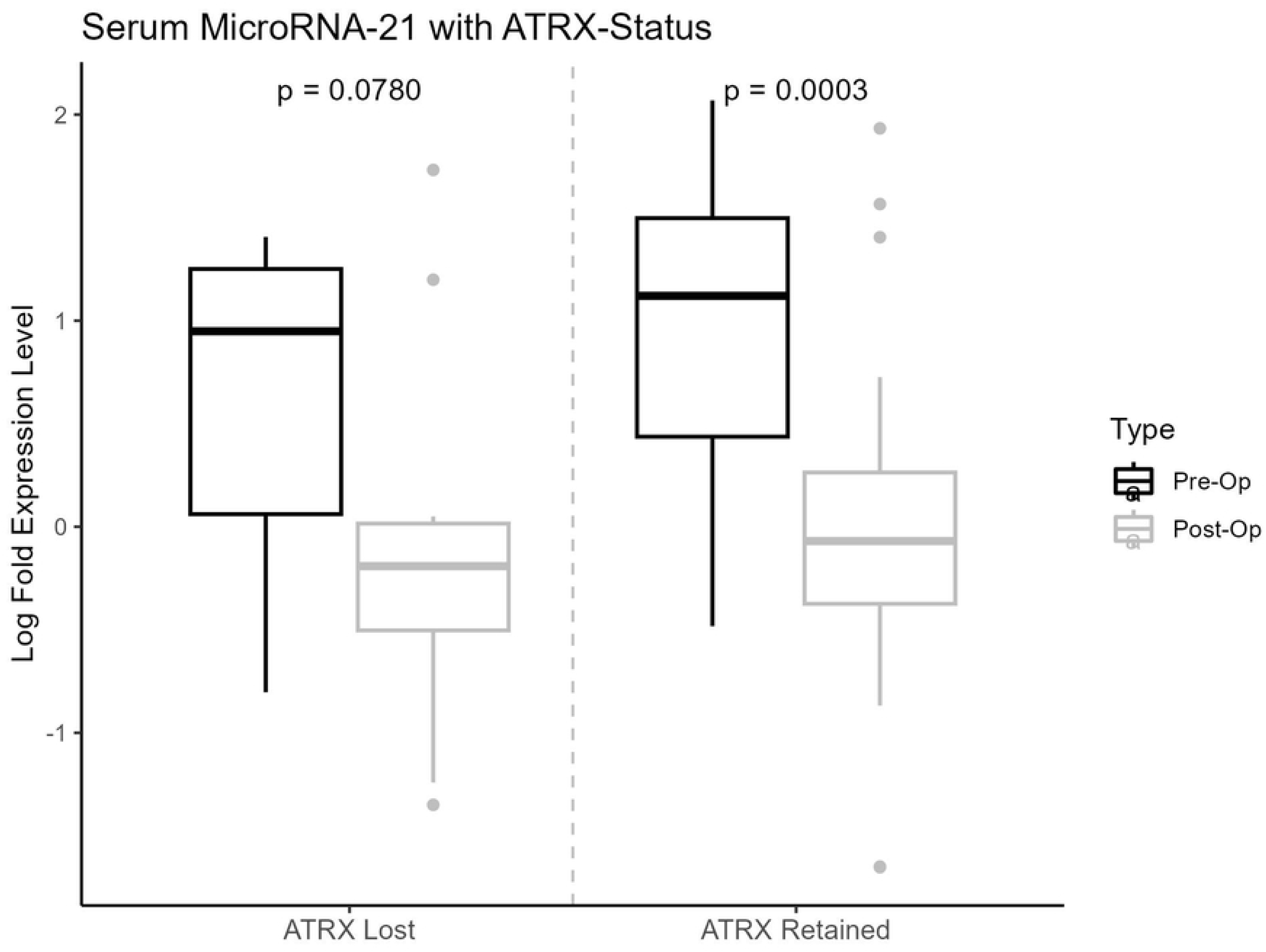
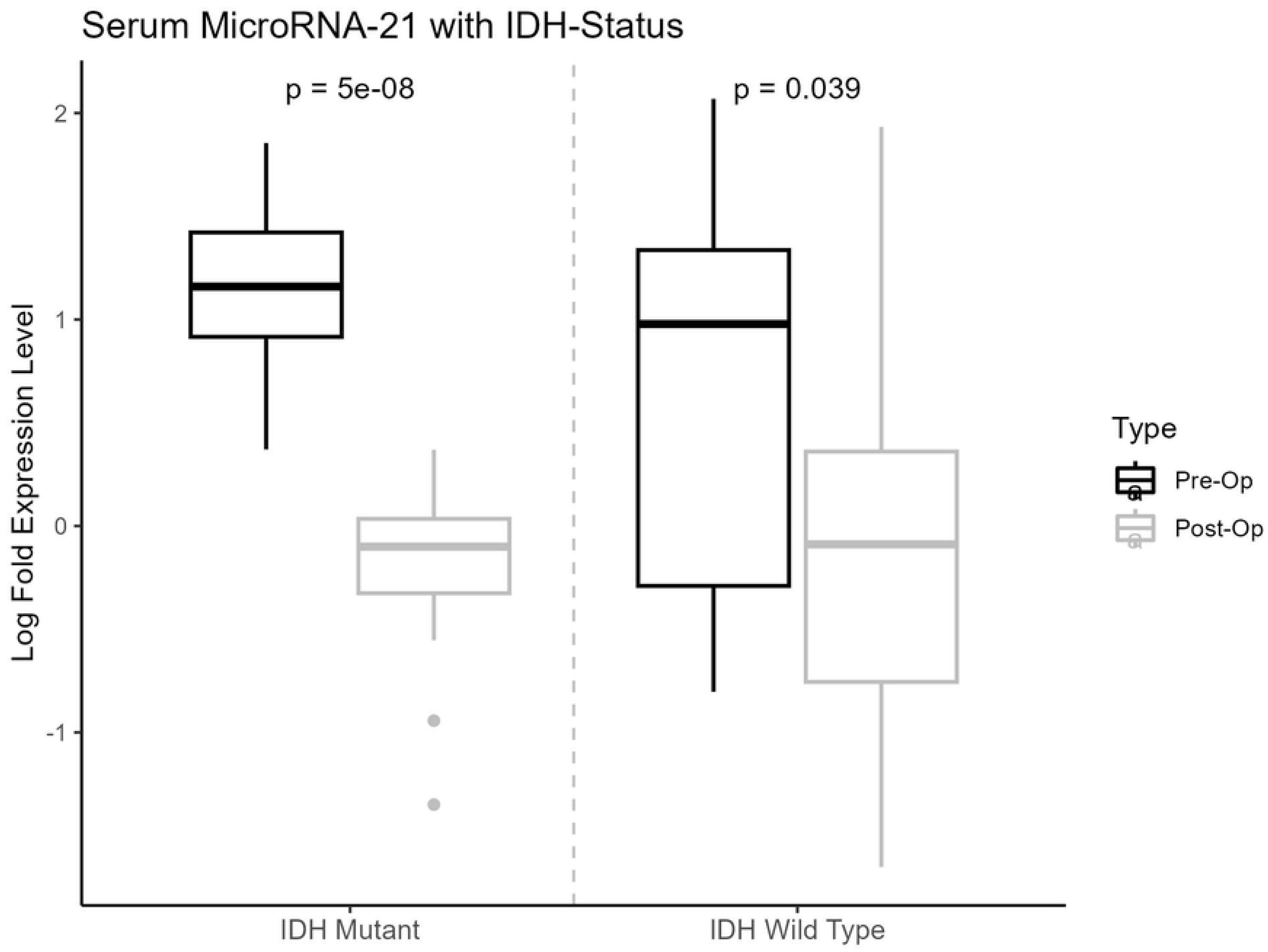
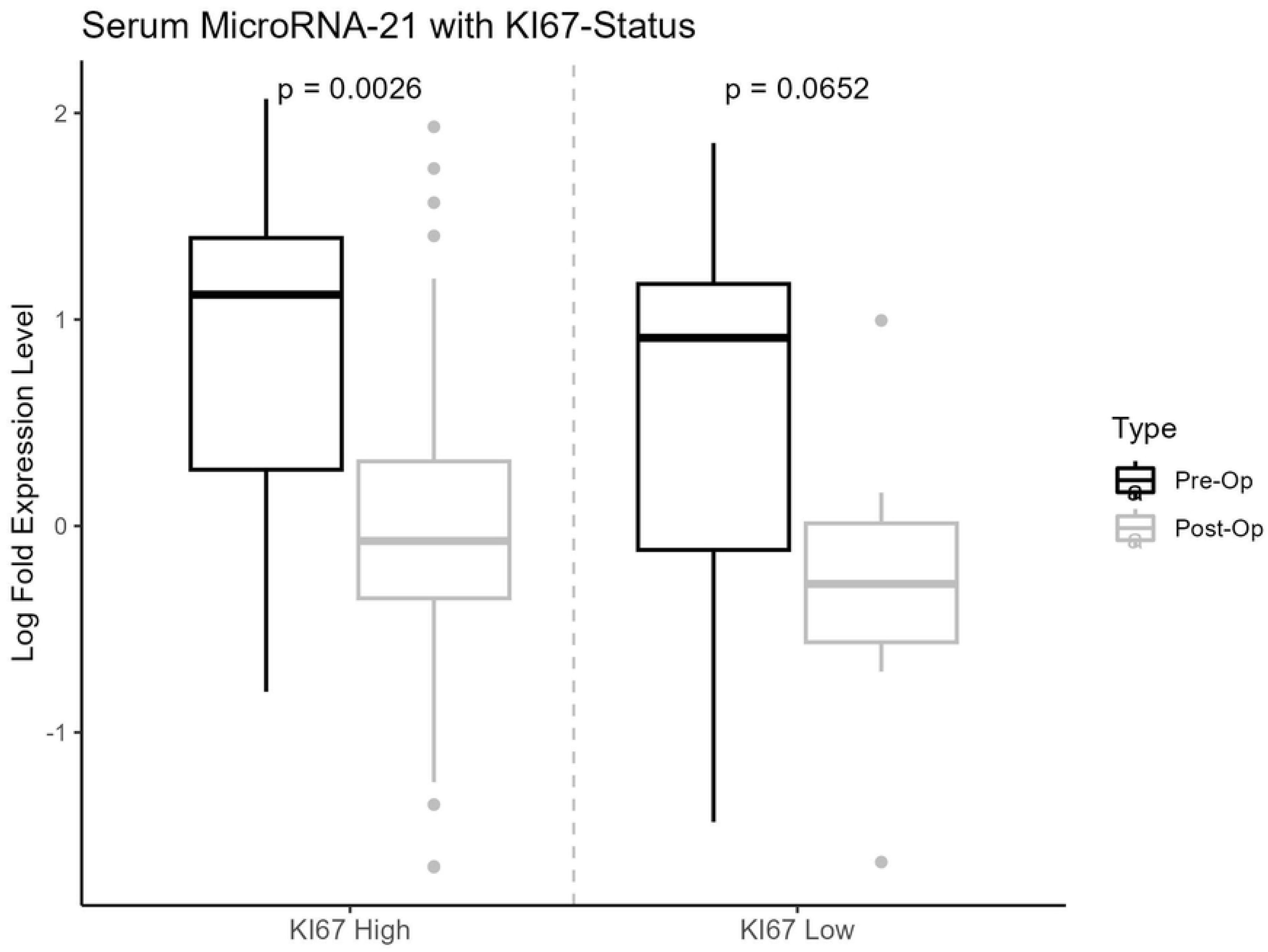
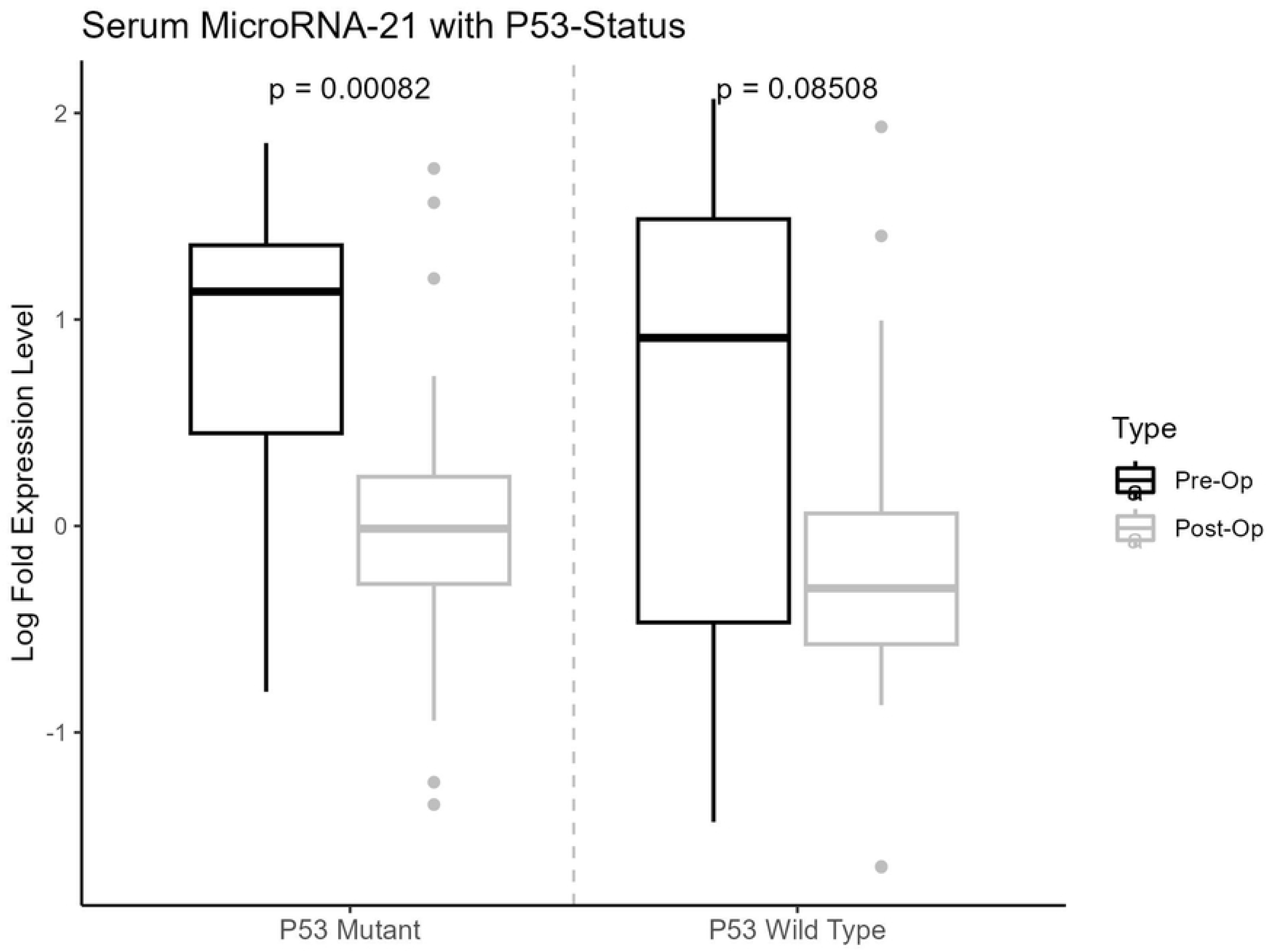
Association of pre- and post-operative serum miR-21 expression with molecular markers, ATRX, IDH, Ki-67 and p53. (Fig.9a) miR-21 expression with ATRX (Lost/Retained). (Fig.9b) miR-21 expression with IDH (Wild Type/Mutant). (Fig.9c) miR-21 expression with Ki-67 (Low/High). (Fig.9d) miR-21 expression with p53 (Wild Type/Mutant).

**Figure 10:**
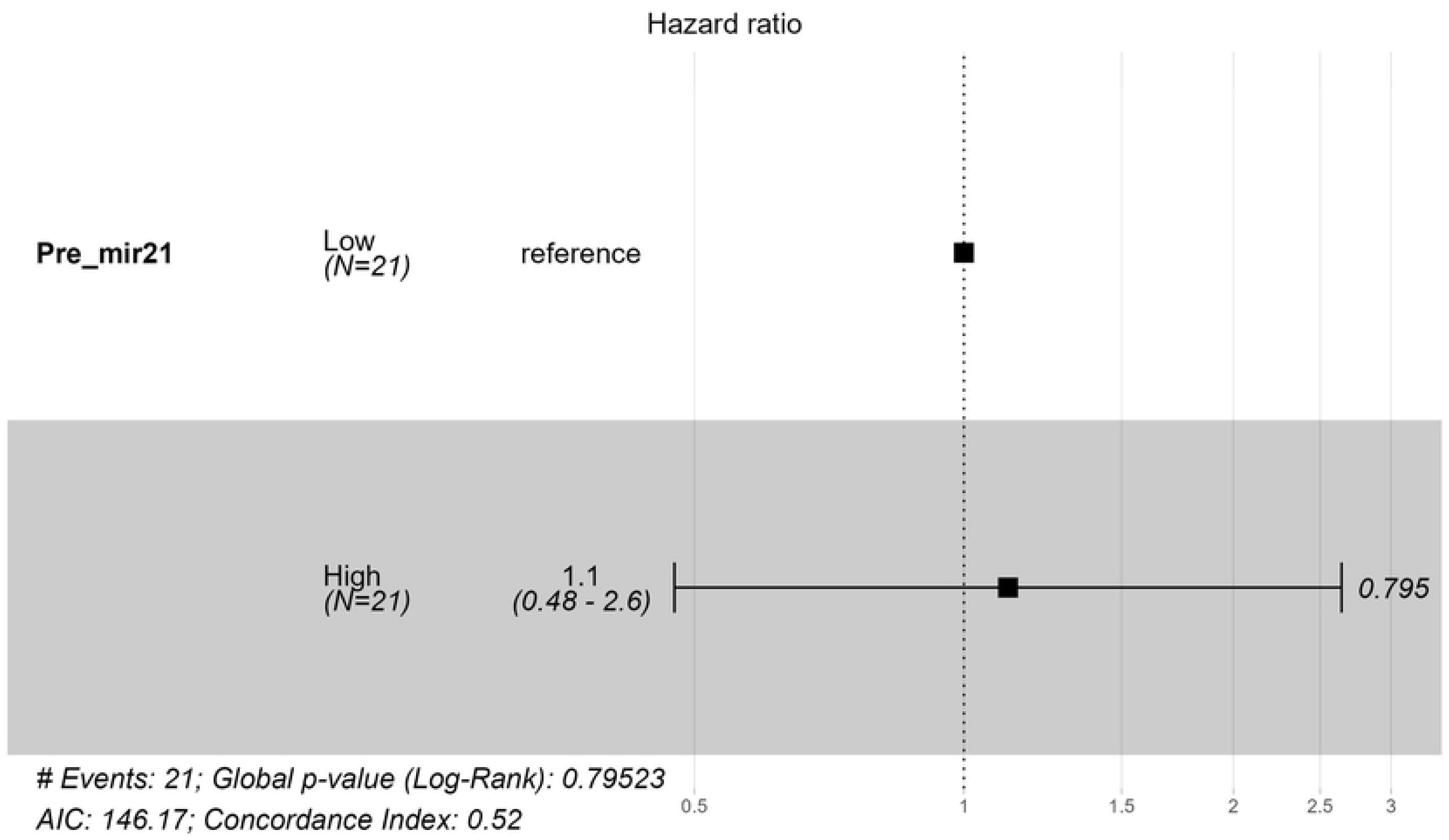
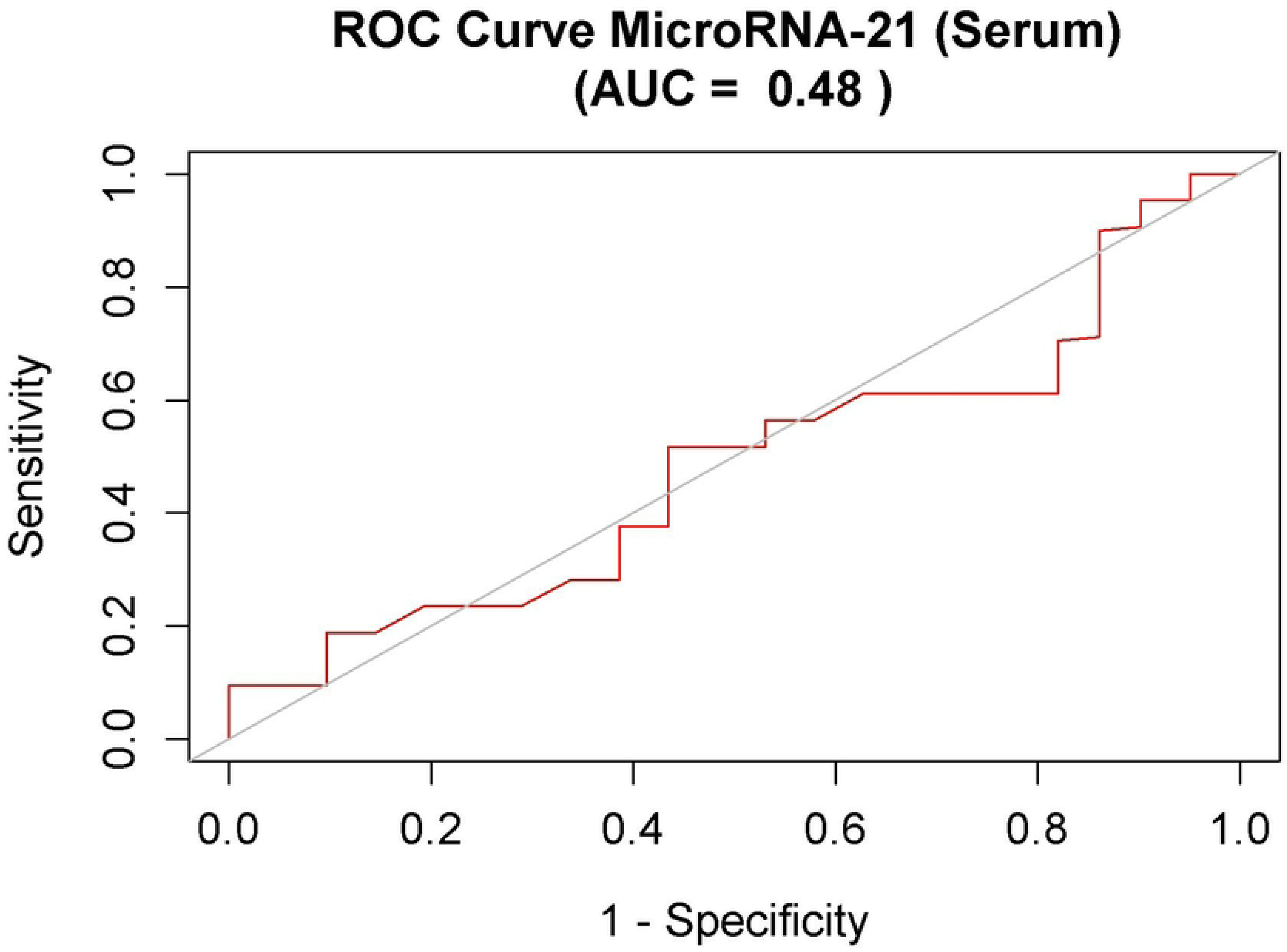
Kaplan-Meier survival curves for high and low expression of miR-21 in pre-operative serum. Overall survival in glioma patient’s preoperative serum with high and low miR-21 expression group.

### Prognostic Value of Serum Mir-21 Expression in Overall Survival

To determine the survival trend Kaplan-Meier survival analysis was done, categorizing the pre-op serum samples into groups of high and low miR-21 expression according to median expression level. The result showed no significant difference between low and high miR-21 expression group with overall survival (p=0.8). Quantitative hazard ratio was also analyzed using Cox regression method in pre-op serum samples from high and low expression groups of miR-21 demonstrating that the group with high miR-21 expression was at 1.1 times higher risk of mortality (95% confidence interval: upper 2.6, lower 0.48) than the patients from low expression group. ROC curve for miR-21 utility in pre-operative serum samples was also plotted which indicates its potential as a useful prognostic marker in the serum of glioma patients (AUC= 0.48).

### Functional and Pathway Enrichment Analysis of Mir-21 Target Genes Using Metascape

We employed the MiRDB database (https://mirdb.org) [16, 17] to identify target genes for miR-21, specifically including genes with a target score of 50 or above in MiRDB [18]. To understand the biological processes involved, we conducted functional enrichment analysis using Metascape (http://metascape.org), focusing on pathways related to cancer, cell cycle and survival. Systematic exploration utilized KEGG pathway (https://www.kegg.jp/en/) [19, 20] WikiPathways, and oncogenic signature gene set enrichment analysis, with an adjusted p < 0.05 as the cutoff value. For miR-21 targeted genes, the analysis uncovered significant involvement in 48 KEGG pathways, 72 Wiki Pathways, 136 Reactome gene sets and 35 Canonical pathways. Notable pathways included were MAPK signaling, cancer pathways, EGF/EGFR signaling, glioma, Glioblastoma signaling, Ras signaling, EGFR tyrosine kinase (Figure 11).

**Figure 11:**
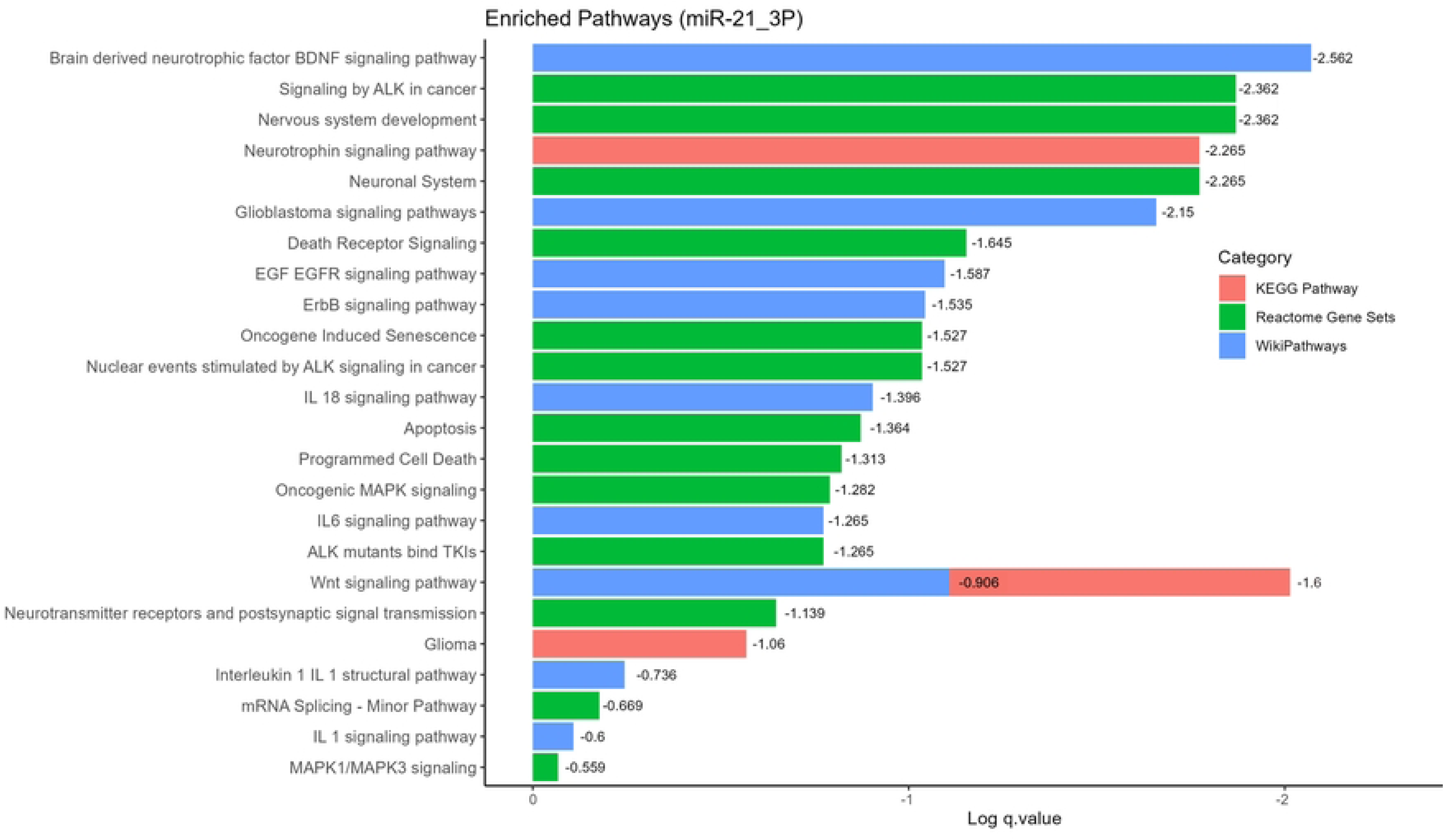
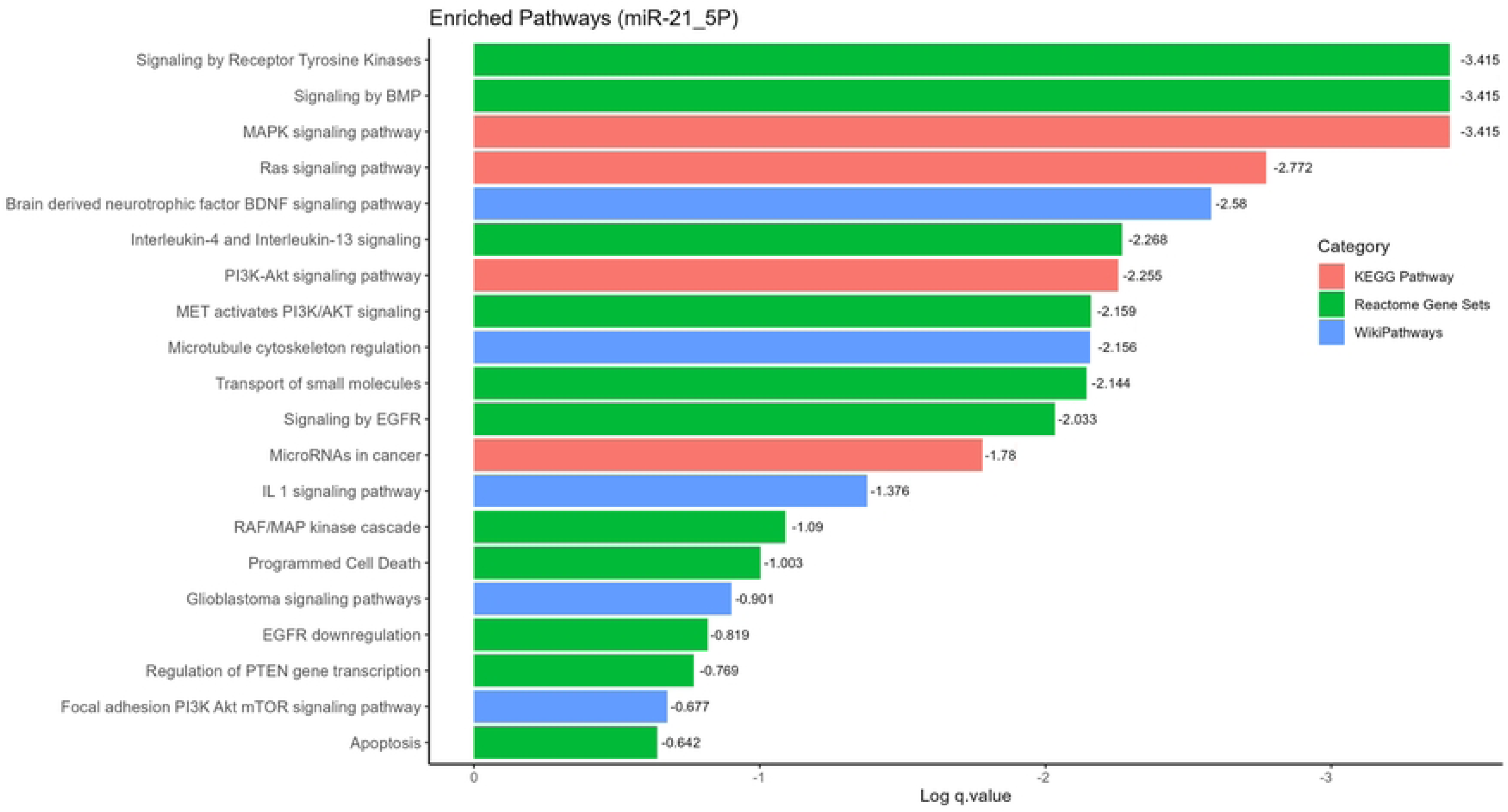
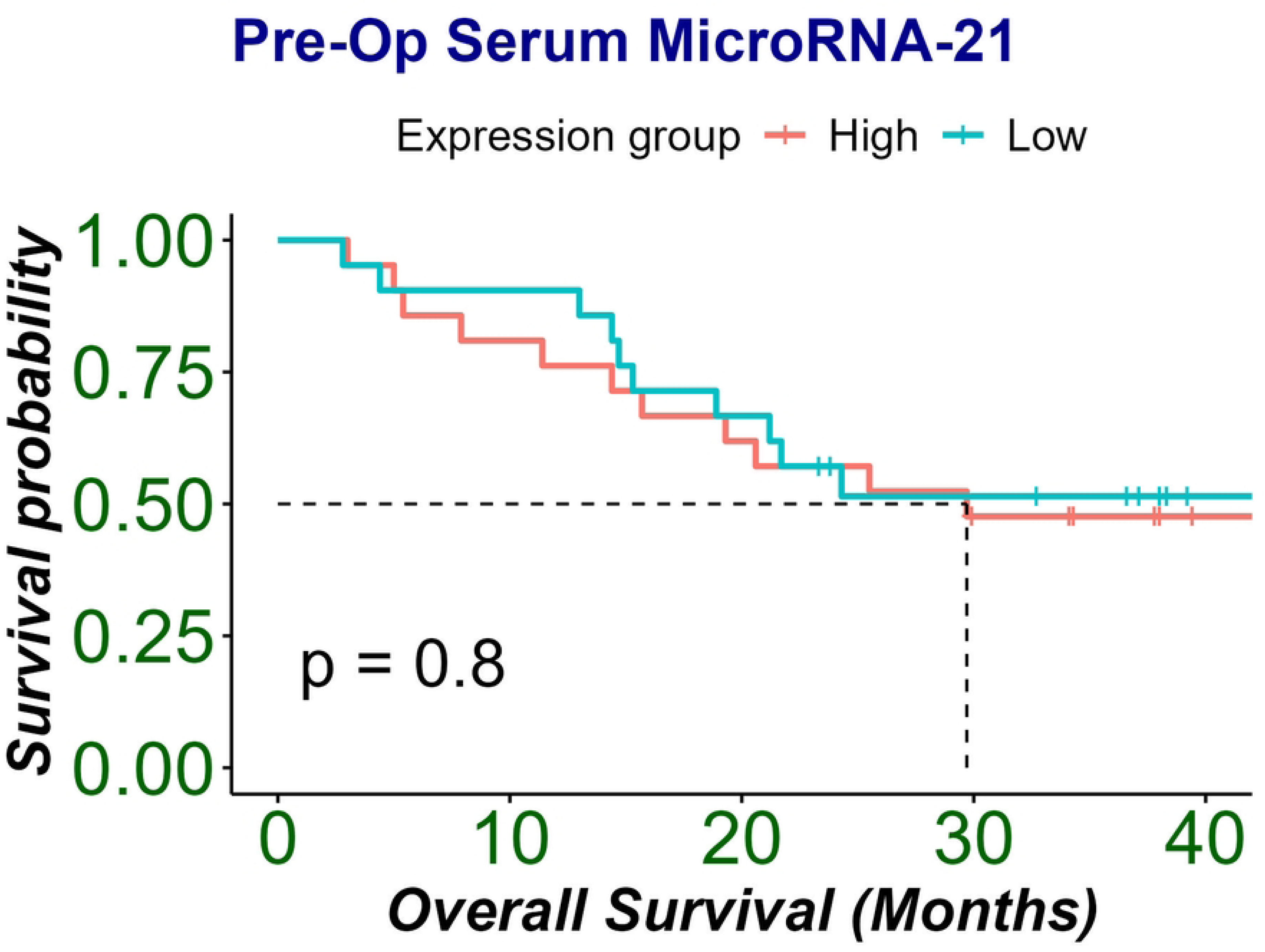
Functional enrichment of miR-21 targets genes. Kyoto Encyclopedia of Genes and Genomes (KEGG) pathways and Wiki Pathways for miR-21(miR-21-3p and miR-21-5p) target genes.

## Discussion

The prognosis of glioma remains a significant challenge due to its complex pathogenesis. Despite advances in treatment strategies and surgical interventions, the diagnosis and overall survival of glioma patients remain poor [21]. The search for non-invasive tumor markers to aid diagnosis is one of the most evolving areas in cancer research. We explored miR-21 as a potential biomarker due to its reported association with GBM aggressiveness in previous studies by various research groups. Our goal was to validate its role specifically in the Pakistani population across different grades of gliomas thus establishing its utility as a non-invasive biomarker. This is especially valuable, as non-invasive biomarkers offer a more accessible and cost-effective alternative to conventional diagnostic approaches, facilitating frequent monitoring of glioma progression and treatment outcomes.

The expression levels of miR-21 were analyzed in patients’ tumor tissues as well as in serum samples. Serum and plasma have been the subject of extensive research for years. However, serum-based tests suitable for widespread use in early tumor detection are still currently limited. Blood samples were collected both before and after the surgery to investigate the correlation between tissue and serum miR-21 expression patterns. Additionally, comparing pre- and post-surgery serum levels allows for evaluating the direct impact of the tumor mass on elevated expression levels and determining whether tumor removal leads to a subsequent decrease in expression.

In this study, we conducted a comparative analysis of miRNA-21 expression in tumor tissue and serum. We collected surgically removed tumor tissue and pre- and post-surgery blood samples from glioma patients, and miR-21 expression levels were analyzed using RT q-PCR. Most patients were male and predominantly under 50 years of age, with significant miR-21 upregulation observed in patients over 50 (p<0.001). There may be a socioeconomic bias in gender and age distribution when analyzing miR-21 expression [22].

miR-21, an oncomiR in glioma, showed significant upregulation in glioma tissues, strongly correlating with tumor characteristics and invasiveness [23]. Similar expression was observed in studies where elevated miR-21 levels were reported with increased cancer grades in breast, esophageal, gastric, colorectal, lung, and epithelial ovarian cancers. miR-21 expression level significantly increased with tumor grade, consistent with previous studies [24],[25]. This increase in miR-21 levels may be linked to its role in angiogenesis and tumor proliferation, key features of grade 4 tumors [26, 27]. No miR-21 expression was detected in adjacent non-malignant brain tissue, indicating its elevated expression was localized to tumor cells [25] Similarly, we found no significant correlation between tumor location and miR-21 expression, which could be due to unavailable patient data.

The latest WHO CNS5 2021 glioma tumor classification emphasizes the use of genetic alterations in IDH, ATRX, Ki-67, and p53 in association with glioma patient overall survival. These markers are significantly associated with disease prognosis and play a crucial role in designing treatment strategies for better glioma outcomes [2]. The significant difference in tissue miR-21 expression among grades (1-4) was validated with previous studies (p <0.005). Additionally, miR-21 expression is related to IDH (mutation/wildtype), ATRX (lost/retained), p53 (mutation/wildtype), and Ki-67 (high/low) index in gliomas, which can help improve cancer prognosis. The results showed a strong link between miR-21 expression and IDH1 mutations. Gliomas with IDH mutations had lower miR-21 expression than IDH wild-type tumors (p=0.03). Furthermore, miR-21 expression and MIB/Ki-67 index demonstrated concurrent changes.

Our results showed that miR-21 expression levels were consistent in the p53 wildtype and mutant groups, suggesting a possible role of MDM2 proteins that work against p53 and suppress its expression at the protein level. Thus, despite the gene expression, the protein is rendered nonfunctional [28],[29]. These findings align with our results and acknowledge that low miR-21 expression is significantly related to better overall survival (OS) in tissue samples of glioma patients [8]. Moreover, study findings distinctively revealed the significant correlation between low miR-21 expression and IDH mutant gliomas, which are associated with a good prognosis of disease. Similarly, significantly better clinical outcomes were found in patients with a low Ki-67 index along with low miR-21 expression in tumor tissues (log-rank test: p= 0.0001).

It is important to note that 76% of patients in the low miR-21 expression group remained alive until the 30^th^ month of survival follow-up, whereas in the high miR-21 expression group, 50% of patients had died by the 20^th^ month of follow-up. Additionally, IDH wildtype patients with low miR-21 expression had a survival probability of 50% at the 30^th^ month, while IDH wildtype patients with high miR-21 expression had a survival probability of only 30% (p<0.0001). This trend in OS of the high miR-21 expression group with wildtype IDH is consistent with previous reports in GBM [30],[31]. A panel comprising molecular markers and miR-21 expression levels analyzed in tissue samples of glioma patients could serve as promising prognostic and predictive biomarkers for glioma. The prediction power was confirmed by ROC analysis with an AUC curve of 0.74. A meta-analysis of nine studies revealed that high miR-21 expression is associated with a poor prognosis in glioma patients [32],[33].

Circulating miR-21 can be detected in glioma patients’ bio-fluids as a non-invasive biomarker. Serum can be a source of microRNAs as they can pass through the blood-brain barrier and remain stable in the blood [33, 34]. This retrospective study’s results add to the current knowledge in the establishment of miR-21 as a circulatory and prognostic marker for early detection of glioma. To determine the difference in serum miR-21 expression before and after surgical removal of the tumor, we analyzed 42 paired serum samples along with 8 healthy controls. miR-21 was found to be differentially expressed in pre-op serum of grade 2, 3 and 4 when compared with controls (p= 0.0031), (p=0.0031) and (p= 0.033) respectively. Moreover, the serum expression levels were also significantly higher in pre-operative glioma samples than in postoperative serum of grade 2 (p=0.0079), grade 3 (p=0.0079) and grade 4 (p=0.0096) which is consistent with previous findings [23, 27]. However, other studies have found a reduction in miR-21 expression post-operation but, the change was not significant, and the author suggested that post-operative sampling should be delayed for more than 24 hours. Since distribution of patients by grade was not uniform in the serum samples in our study, with grade 4 comprising (61.90 %), grade 3 (11.90%), grade 2 (11.90%) and grade 1 (14.28%) of the sample, this skewed the median distribution of the miR-21 expression group. Consequently, the study result may partly reflect a high miR-21 expression in low grade groups and our result showed no significant difference between low and high miR-21 expression groups in terms of overall survival (p=0.8), with an AUC curve of 0.48.

The discrepancy between survival analyses of tissue and serum samples can be attributed to several key factors. First, the serum sample cohort (n=42) was less than half the size of the tissue sample group (n=90), which reduces statistical power to detect survival differences. Second, there was a notable difference in grade distribution between the cohorts. The serum sample cohort had a substantially higher proportion of grade 4 gliomas (61.90% vs 48.8% in tissue samples) and a lower proportion of grade 2 tumors (11.90% vs 26.6% in tissue samples). Given that we stratified patients into high and low expression groups based on median miR-21 expression, this skewed grade distribution likely resulted in grade 4 patients being distributed across both expression groups in the serum cohort. Since grade 4 gliomas are associated with poorer survival outcomes, their presence in both high and low expression groups may have masked any survival differences that could be attributed to miR-21 expression levels alone. This is supported by the hazard ratio analysis, which showed a much stronger prognostic value in tissue samples (HR: 3.4, p<0.001) compared to serum samples (HR: 1.1, 95% CI: 0.48-2.6).

Enrichment analysis has also provided molecular insights into signaling pathways and transcription factors (TFs) regulated by miR-21, which could serve as therapeutic targets in glioma prognostics. miR-21, which has been established as an onco-miR, possesses oncogenic potential involved in cell proliferation, differentiation, apoptosis, and radio/chemosensitivity. It is reported to be overexpressed in several malignancies, including glioma. Our function enrichment analysis revealed the involvement of miR-21 in Ras signaling, NTRK, receptor tyrosine kinase, PI3/AKT/MET, IL6, and FGF pathways. These pathways are well known to be linked with GBM pathogenesis [34].

PIK3R1 is directly targeted and negatively regulated by miR-21. Studies have shown that PIK3R1, coding for the protein p85α, acts as a tumor suppressor by inhibiting the PI3K/AKT signaling pathway, which is known for promoting cell proliferation, migration, and invasion in cancer [35]. When miR-21 is overexpressed, it leads to the downregulation of PIK3R1, resulting in more aggressive tumor behavior, this relationship is further evidenced by miR-21 knockout models, where the removal of miR-21 restores PIK3R1 activity, leading to reduced PI3K/AKT pathway signaling and, consequently, the inhibition of tumor growth and invasion [36]. PI3K/AKT/mTOR signaling pathway is also regulated by miR-21-5p and this pathway is involved in autophagy regulation in gliomas. Other pathways known to be involved in miR-21-assisted glioma progression include Ras/MAPK signaling. These pathways are found to be abnormally activated in gliomas. Studies have found that miR-21 targets Spry2, a negative feedback regulator of Ras, and consequently amplifies Ras/MAPK signaling particularly in conjunction with dysfunctional PTEN. This results in proliferation, migration, invasion, and metastasis of cancer cells. Therefore, miR-21 overexpression may promote glioma growth via overactivation of Ras/MAPK [37].

Another important gene target of miR-21 is the FOXO gene family. These genes play a key role in gluconeogenesis and glycogenolysis regulation. Studies have revealed that FOXO family genes could be tumor-inhibiting. Here, miR-21 acts as an oncogene, binding to FOXO1 transcripts and terminating its translation. This mechanism could also account for the association of poor prognosis in glioma with miR-21 overexpression by inducing chemoresistance in tumor cells [38].

Several important candidate genes have been identified as miR-21 targets, amongst which FASLG is the most intriguing. FASLG and its receptor FAS have been regarded as crucial effectors of apoptosis in various biological processes. Studies have linked their downregulation in many carcinomas, highlighting their tumor suppressor role by regulating cellular apoptosis and proliferation. miR-21 is involved in regulating apoptosis and growth of Glial Stem Cells (GSCs) through downregulating FASLG proteins. Thus, FASLG serves as a direct miR-21 target in GSCs regulating its proliferation [39]. GAB1 a direct target of miR-21, is an important gene for tumor angiogenesis and metastasis, it is also involved in diverse biological processes such as cellular proliferation and differentiation, survival, angiogenesis, and inflammation. GAB1 allows interconnection of different cellular signaling pathways, notably EGFR/PI3K/AKT pathway. It could also regulate MAPK and ERK pathways resulting in a positive feedback loop and conversely EGF stimulation could induce GAB1 phosphorylation inhibiting cell signaling via PI3K/AKT pathway [40]. PTEN acts as a tumor suppressor through inhibiting PI3K/AKT pathway, is a well-established miR-21 target. miR-21 is associated with autophagy in glioma through the PI3K/AKT/mTOR pathway, and its high expression is linked to high-grade glioma [41]. FOXO, a transcription factor for FASL, functions as tumor suppressor and a downstream molecule of PI3K/AKT pathway. FOXO1 is phosphorylated by activated AKT and is degraded by proteosomes in cytoplasm. It is hypothesized that miR-21 might modulate the PI3K/AKT pathway by targeting PTEN upstream and FOXO1 downstream. In glioma, STAT3 signaling pathway genes, and its upstream genes can be targeted by miR-21. STAT3 binds to miR-21 promoter and regulates its expression. miR-21 also regulates STAT3 gene activation and expression of hTERT in a STAT3-dependent manner [42].

The p53 tumor suppressor gene is the hub of cellular pathways responding to DNA damage, aberrant mitogenic stimulations and cellular stress. P53 is activated by these signals and facilitates growth arrest, promotes apoptosis or can mediate DNA repair mechanisms. The presence of mutations in p53 in almost all cancer types describes its tumor preventive role. Studies show that p53 is activated by factors responding to oncogenic stimuli and assist p53 by either stabilizing it or acting as cofactors. These co factors could either start transcription activation or repression of genes to promote cell cycle arrest or apoptosis. These cofactors are transcription factors like TP53BP2, TOPORS, DAXX, TGF-β and many others [28]. TGF-β, is the quintessential growth inhibitory cytokine that can also induce apoptosis. Computational studies have revealed correlation between miR-21 and components of TGF-β pathway in underlying carcinogenesis [11]. Another important transcription factor that plays a key role in tumor proliferation, angiogenesis and invasion is STAT3, which is a member of STAT family TFs. It is a critical inducer of mesenchymal transformation in glioma. A study has reported that STAT3 and miR-21 interact closely and form a regulatory loop. Also, studies suggest lower levels of STAT3 after miR-21 inhibition in cell treatments. This provides evidence that miRNA-21 provides some regulatory feedback to STAT3.

miR-21 can be activated by a variety of growth factors and cytokines namely, EGFR, IL-6R, JAK and other kinases. The IL-6/STAT3 signaling axis holds great importance in glioma, which also explains the overexpression of miR-21 through the induction of STAT3. IL-6 is a prime tumor promoting factor secreted by malignant cells in tumor microenvironment. It stimulates proliferation and cancer development. The increase of IL-6 the master TF, STAT3 translocate to nucleus and induces miR-21 through binding to its upstream enhancer region. After which miR-21 enhances NF-κB and IL-6 upregulation. This upregulation of NF-κB and IL-6 induces miR-21 expression and forms a positive feedback loop in response to any inflammatory stimulus or DNA damage [43]. In certain cancers the activation of MAPK signaling targets miR-21 overexpression by KRAS, leading to oncogenesis. It induces cell migration, invasion, and drug resistance through various tumor suppressors’ inhibition. This increased expression of miR-21 targets TGF-β-activated SMAD-3/4 leading to epithelial-to-mesenchymal transition (EMT) [29]. Antiapoptotic miRs target apoptotic genes that are usually overexpressed in GBM. miR-21 is an anti-apoptotic that promotes tumor promotion by targeting cell signaling pathways of P53, TGF-β, mitochondrial apoptotic pathways. Further, it also modulates extrinsic apoptotic pathway through FASL downregulation. This ability of mir-21 is evident in cancer stem cells, making it a critical player in GBM pathogenesis and a promising target for therapeutic interventions.

This study has some limitations, including the relatively small sample size and the fact that it is a single-center study. Future research should focus on larger, multi-center studies to validate these findings further and explore the mechanisms by which miR-21 influences glioma progression and response to treatment.

## Conclusion

Our findings suggest that miR-21 is significantly upregulated in glioma tissues and serum of glioma patients, correlating with tumor grade, molecular markers, and overall survival. It may serve as a non-invasive biomarker for glioma prognosis and monitoring. Further research is needed to establish miR-21’s clinical utility in a broader context and to explore its potential as a therapeutic target.

## Supporting Information

### Contributions

Conceptualization: NM

Methodology: WA, SN, SU, AAL

Data Curation: WA, SN, AAL

Supervision: NM

Formal Analysis: SS

Original Draft Preparation: NM, WA, UA, SS, SI, AAL

Review & Editing: NM, SHA, SAE, SI, AAL

### Ethics declaration

The authors declare no competing financial and non-financial interests

### Funding

This study was funded by the Higher Education Commission (Ref No. 20-16919/NRPU/R& D/HEC/2021)

### Informed Consent Statement

Written informed consent has been obtained from all participants to publish this paper

